# Daily fluctuations in spinal adenosine determine mechanisms of respiratory motor plasticity

**DOI:** 10.1101/2022.12.15.520642

**Authors:** Alexandria B. Marciante, Yasin B. Seven, Mia N. Kelly, Raphael R. Perim, Gordon S. Mitchell

## Abstract

Plasticity is a fundamental property of the neuromotor system controlling breathing. One key example of respiratory motor plasticity is phrenic long-term facilitation (pLTF), a persistent increase in phrenic nerve activity after exposure to intermittent low oxygen or acute intermittent hypoxia (AIH). pLTF can arise from distinct intracellular signaling cascades initiated by serotonin and adenosine; these cascades interact via powerful crosstalk inhibition. We demonstrate the serotonin/adenosine balance varies dramatically with time-of-day and details of the AIH protocol. Using a “standard” AIH protocol, the mechanism driving pLTF shifts from serotonin-dominant, adenosine-constrained during rest, to adenosine-dominant, serotonin-constrained in the active phase. This mechanistic ‘flip’ results from daily changes in basal spinal adenosine levels across time-of-day combined with hypoxia-evoked spinal adenosine release. Since AIH is emerging as a promising therapeutic modality to restore respiratory (and non-respiratory) movements in people with spinal injury or ALS, new knowledge that time-of-day and protocol details impact mechanisms driving pLTF has experimental, biological and translational implications.

## INTRODUCTION

Neuroplasticity is a fundamental property of neural systems, including the neural system controlling breathing ^1^. One important model of respiratory motor plasticity, known as phrenic long-term facilitation (pLTF), is elicited by repeated exposure to brief episodes of low oxygen, or acute intermittent hypoxia (AIH). pLTF is expressed as a persistent increase in phrenic nerve activity lasting hours after AIH has ended ^2^. AIH also elicits plasticity in other respiratory (*e.g*. inspiratory intercostal, hypoglossal & laryngeal) and non-respiratory motor systems (*e.g*. hand/arm and leg/walking function) ^2,3^. Beyond it’s physiological relevance, repetitive AIH exposure is emerging as a promising therapeutic modality to improve breathing, walking and arm/hand function in people with chronic spinal cord injury ^1,4,5^ and ALS ^6^. However, greater understanding of mechanisms underlying AIH-induced motor plasticity is essential to optimize its application for therapeutic benefit ^3^.

Phrenic LTF was first described as a persistent increase in phrenic nerve activity lasting hours following episodic electrical stimulation of carotid chemo-afferent neurons ^7–9^ or 3, 5 minute episodes of moderate hypoxia ^8,10,11^. With electrical chemo-afferent neuron activation ^12^ and moderate AIH ^13^, pLTF is dominated by a serotonin-dependent mechanism involving cervical spinal serotonin 2 (5-HT_2_) receptor activation ^14,15^. Conversely, when AIH consists of *severe* hypoxic episodes, pLTF arises from a distinct mechanism that requires cervical spinal adenosine 2A (A_2A_) receptor activation ^16,17^. 5-HT_2_ (G_q_-coupled) *versus* A_2A_ (G_s_-coupled) metabotropic receptors initiate completely distinct intra-cellular signaling cascades, known as the Q and the S pathways to phrenic motor facilitation, respectively ^2,18,19^ (**Figure 1**). The Q and S pathways interact via powerful crosstalk inhibition: when both are equally activated, plasticity is cancelled ^2,20^. Thus, AIH-induced pLTF is not observed when the serotonin and adenosine-dependent mechanisms are activated concurrently by: 1) hypoxic episodes of intermediate severity ^21^; or 2) sustained moderate hypoxia ^22^. With moderate AIH (mAIH), A_2A_ receptor activation is sufficient to constrain, but not eliminate pLTF ^16,23^. Thus, the balance of cervical spinal serotonin versus adenosine receptor activation is a powerful regulator of pLTF.

**Figure 1.**
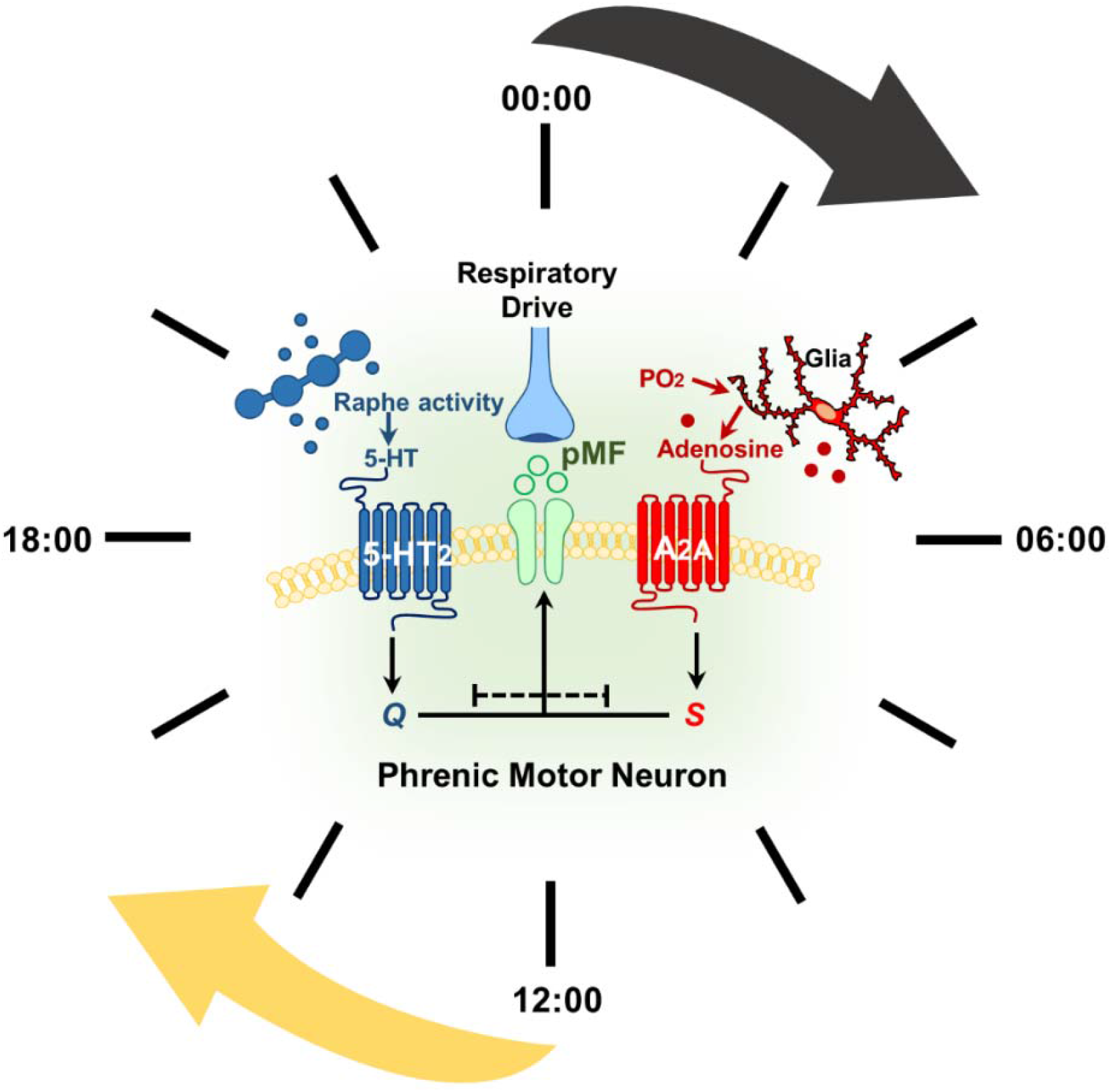
Competing serotonin and adenosine-dependent pathways to phrenic motor facilitation (pMF) interact via powerful crosstalk inhibition. With moderate AIH (mAIH), carotid chemoreceptors are stimulated, increasing breathing and (indirectly) brainstem raphe serotonergic neuron activity. Serotonin release in the phrenic motor nucleus initiates the Q pathway to pMF (blue). However, mAIH also elicits spinal tissue hypoxia, triggering glial ATP release and extracellular adenosine accumulation, which undermines serotonin-dominant plasticity due to adenosine 2A (A2A) receptor activation on phrenic motor neurons (i.e., S pathway; red). Time of day and hypoxic episode duration effects on pLTF and its underlying mechanism(s) are unknown.

Given the important balance between serotonin versus adenosine receptor activation during hypoxia in regulating phrenic motor plasticity, we wondered if physiological conditions known to influence basal adenosine levels in the central nervous system (CNS) impact this relationship and, thus, the magnitude of phrenic motor plasticity. One factor known to impact basal adenosine levels in the CNS is time-of-day, particularly when comparing active *versus* resting phases ^24–26^. Since studies on spinal respiratory motor plasticity have invariably been performed during the day, which is the rest phase in rodents ^2,11^ but the active phase in humans ^27,28^, it is essential to understand the impact of AIH delivered during the daily rest *versus* active phase. Further, since distinct AIH protocols routinely used in rodents (3, 5 minute episodes) *versus* humans (15, 1 minute episodes) may differentially impact hypoxia-evoked adenosine release ^2^ details of the AIH protocol used may also impact respiratory motor plasticity in the rest versus active phases of the day.

During hypoxia, spinal adenosine accumulation arises from glial ATP release, with subsequent conversion to adenosine via the ectonucleotidases ^29,30^. The extent of extracellular adenosine accumulation during hypoxic episodes and the drive to plasticity exerted by spinal adenosine receptor activation depends on the severity and duration of hypoxia ^16,22^. Historically, pLTF is typically studied during the rest phase (day) in rodents, when mAIH generates enough spinal adenosine to constrain serotonin-dominant phrenic motor plasticity ^23^. On the other hand, the same AIH protocol with more severe hypoxic episodes elicits greater spinal adenosine accumulation, shifting pLTF from serotonin to adenosine-dominance ^16^. These “dueling pathways” have profound implications concerning pLTF regulation across the daily rest/active cycle.

Extracellular adenosine levels fluctuate in the CNS across the circadian rest/active cycle ^24,25,29,31^. CNS adenosine levels increase during wakefulness (i.e. active phase), providing a powerful drive to sleep ^25^. Conversely, adenosine decreases during the rest phase ^24,25,32^, minimizing its potential for contributions to respiratory motor plasticity. It is not known if adenosine exhibits daily fluctuations in the spinal cord.

In other regions of the CNS, daily rhythms influence neuroplasticity ^33^. For example, hippocampal, cortical and thalamic synaptic plasticity are all modified across the daily rest/active cycle, reflecting circadian rhythms and/or activity state-dependent changes in extracellular adenosine levels ^31,33–36^. Indeed, daily fluctuations in adenosine modulate synaptic transmission and activity-dependent long-term potentiation in the hippocampus ^33,37,38^. Although hippocampal synaptic plasticity is transient during the active phase since adenosine disrupts LTP consolidation ^39^, plasticity is consolidated and lasts longer during the rest phase due to lower CNS adenosine levels ^34^. There are no reports concerning how the rest/active cycle and associated fluctuations in spinal adenosine level affect AIH-induced pLTF.

Thus, we performed experiments to determine the impact of time-of-day on AIH-induced pLTF. We used a standard experimental preparation because of known advantages and the extensive background information available: anesthetized, paralyzed and ventilated Sprague Dawley rats. Using this experimental preparation, we tested the hypothesis that serotonin-dependent plasticity is undermined in the active phase due to greater basal adenosine levels. We report profound time-of-day effects on the magnitude and mechanism plasticity due to shifts in the serotonin/adenosine balance of the ventral cervical spinal cord due to: 1) fluctuations in ventral cervical spinal cord adenosine levels across the day/night cycle, and 2) the duration/pattern of hypoxic episodes within the AIH protocol. Surprisingly, when using our “standard” rodent AIH protocol (3, 5 min episodes of moderate hypoxia, 5 min intervals), time-of-day completely shifts the mechanism of pLTF from serotonin-dominance in the rest phase to adenosine-dominance in the active phase. However, the duration of hypoxic episodes within the AIH protocol also impacts pLTF expression and mechanism; the same cumulative duration of hypoxia presented as shorter hypoxic episodes (15, 1 min hypoxic episodes, 1 min intervals) greatly augments pLTF during the rest phase, but suppresses pLTF during the active phase. These findings have profound implications with respect to the design of future experiments, our understanding concerning mechanisms and functional significance of phrenic motor plasticity, and on our efforts to harness AIH-induced motor plasticity to treat humans suffering from chronic spinal cord injury or other neuromuscular disorders that compromise breathing and other movements.

## RESULTS

### Basal ventral spinal adenosine fluctuates across the daily rest/active cycle

While adenosine levels fluctuate throughout the daily rest/active cycle in multiple brain regions ^24,32^, it is unknown if the spinal cord exhibits a similar pattern. There is evidence that circadian clock genes and genes associated with serotonin-dependent phrenic motor plasticity oscillate within the phrenic motor system ^40^, providing support for the potential presence of spinal adenosine fluctuations across the rest/active cycle. To test for daily adenosine fluctuations, ventral cervical segments 3-5 (i.e., segments containing the phrenic motor nucleus) were harvested in mid-rest (12 PM) and mid-active (12 AM) phases to assess basal adenosine and serotonin levels (n = 6 per group). Care was taken to minimize light cues ^41,42^; tissues collected in the midactive phase were harvested under transient dim red light. Spinal adenosine levels were measured with a fluorometric assay (Cell BioLabs), adapted from Jagannath et al., 2021 ^41^. Ventral spinal C_3_-C_5_ adenosine was significantly higher during the midactive (14.5 ± 1.8 μM) *versus* the midrest phase (7.1 ± 1.4 μM; t_10_ = 3.099, p = 0.011; unpaired *t*-test; **Figure 2A**). Given the importance of serotonin in AIH-induced pLTF, basal serotonin levels were also measured using a rat serotonin ELISA (MyBioSource). There was no difference in serotonin levels between midrest (46.6 ± 3.3 ng/mL) versus midactive (46.3 ± 2.3 ng/mL) (t_11_ = 0.084, p = 0.935; unpaired *t*-test; **Figure 2B**), consistent with a prior report in the rat lumbar spinal cord ^43^.

**Figure 2.**
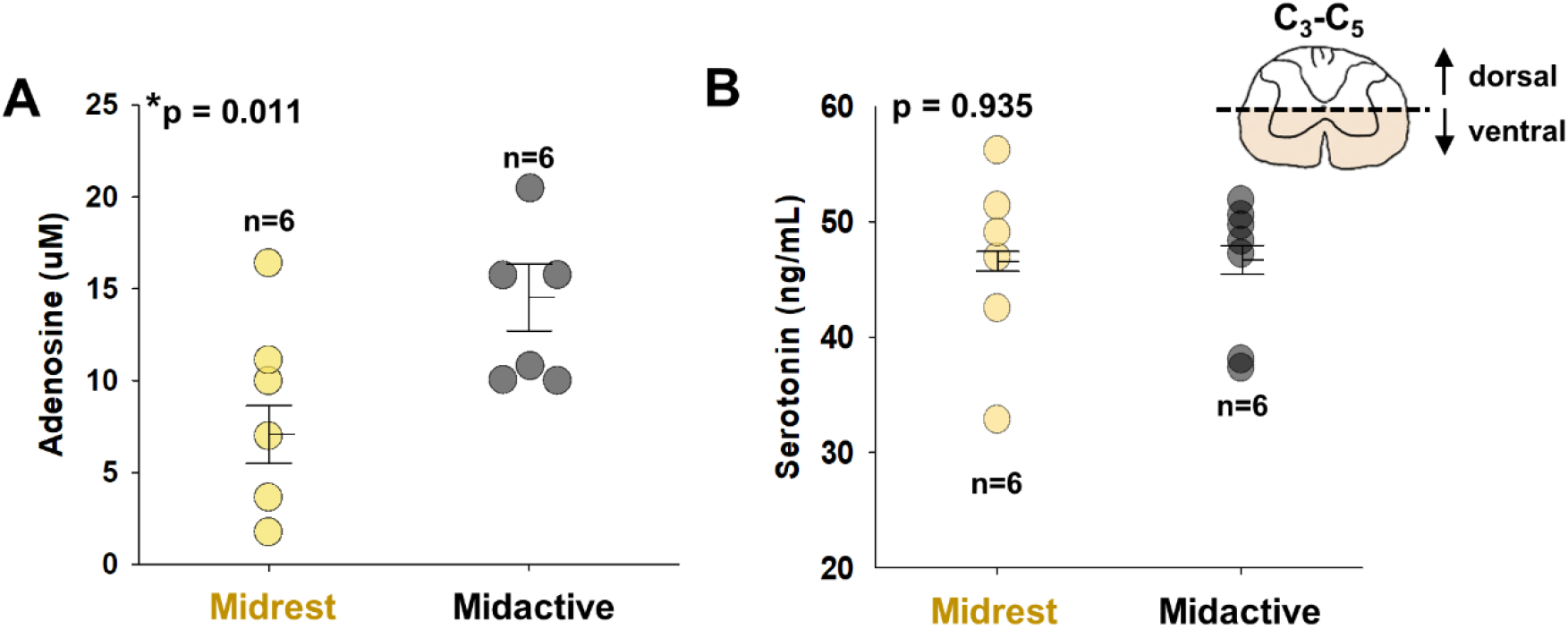
Basal spinal adenosine levels are greater in active *versus* rest phase. **A**: Ventral spinal C_3_-C_5_ adenosine concentration is significantly elevated during midactive (midnight; black) *versus* midrest (noon; yellow) phase (p = 0.011). **B**: Basal serotonin levels measured at these same time points are not different (p = 0.935). All groups, n = 6; comparisons were made using unpaired *t*-test. Bars show means ± 1 SEM; p < 0.050.

We also measured adenosine 2A (A_2A_) receptor protein levels (**Figure S1**) in the same region, and found no differences between midrest and midactive samples; previous work indicates no difference in ventral spinal 5-HT_2A_ receptor expression at either time ^40^. Thus, ventral cervical spinal adenosine levels fluctuate across the daily rest/active phases in spinal regions associated with the phrenic motor nucleus that may regulate mAIH-induced pLTF across the day/night cycle.

### pLTF magnitude is the same at noon and midnight when using the “standard” moderate AIH protocol

Moderate AIH (3, 5 min hypoxic episodes, 5 min duration; arterial PO_2_ > 40 mmHg) has been used in most studies of pLTF ^3^. Thus, we recorded phrenic nerve activity in anesthetized, paralyzed, vagotomized and artificially ventilated rats prepared for phrenic nerve recordings (for details concerning experimental preparation, see *Methods* or: ^15,44^). In the midactive phase, care was taken to prevent light cues—rats were prepared for surgery and anesthetized under a dim red light before being transferred to the experimental rig. After transferring rats to the rig, their eyes were immediately covered. Once adequate anesthesia was confirmed by lack of a toe-pinch reflex, room lights were turned on, but their eyes remained covered for the duration of experiments.

The ventilator tidal volume (mL) was set for each rat based on body mass (0.007 mL * body mass, g). Ventilator rate was maintained between 72-74 breaths per minute. During baseline conditions, rats were ventilated with 60% O_2_ (balance N_2_). Inspired CO_2_ was adjusted during surgery to maintain end-tidal PCO_2_ between 38-41 mmHg. Since CO_2_ influences phrenic nerve burst amplitude and frequency ^8,9,13,45,46^, apneic and recruitment CO_2_-thresholds were determined and used as a guide to standardize the arterial partial pressure of carbon dioxide (PaCO_2_) during baseline conditions (recruitment threshold + 2 mmHg PCO_2_) and was carefully maintained within ±1.5 mmHg of baseline throughout the experiment (**Table S1**). Rectal temperature was measured and maintained at 37.5 ± 1.0°C.

The standard mAIH protocol consisted of 3, 5 min episodes of moderate hypoxia (arterial partial pressure of oxygen, PaO_2_: 40-55 mmHg), with 5 min intervals of baseline O_2_ ^8,10,11^ (**Figure 3A**). Following mAIH, phrenic nerve activity was measured for 90 minutes after baseline conditions were restored (PaO_2_ > 150 mmHg; **Figures 3A** and **3B**). Moderate AIH was delivered during midrest (12 PM) or midactive (12 AM) phases. Blood gas measurements were taken 2-3 times during the initial baseline, the last minute of hypoxic episodes, and at 30, 60, and 90 min post-mAIH (**Tables S1** and **S2**). One-minute averages of phrenic nerve amplitude were measured 90 min post-mAIH, and are presented as percent change from pre-AIH baseline. Raw integrated phrenic nerve amplitude at baseline and during maximal chemoreceptor stimulation (10% O_2_, 7% CO_2_, balance N_2_) delivered at the end of each experiment are included in **Table S3** to assess recording quality. Respiratory frequency is shown in **Table S4**.

**Figure 3.**
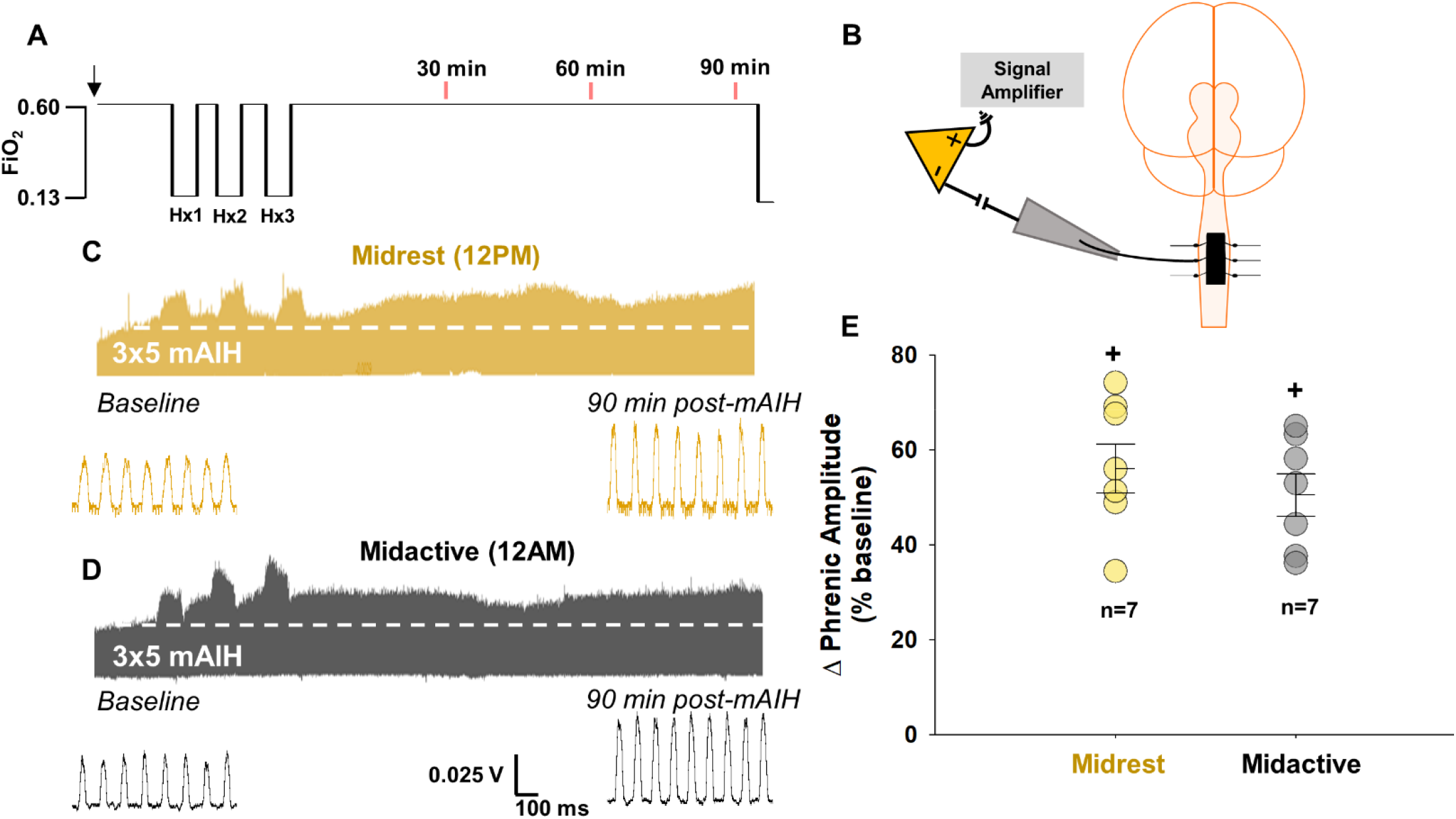
Phrenic LTF elicited by 3×5 mAIH is similar during midrest and midactive phases. 3×5 mAIH was delivered at noon or midnight to rats on a normal light cycle (**A**). Phrenic nerve activity was recorded in urethane anesthetized rats before, during and after 3×5 mAIH (**B**). Representative compressed neurogram of integrated phrenic nerve activity during baseline, mAIH (Hx1-3; PaO_2_ = 40-55 mmHg), and for 90 min after restoring baseline conditions during midrest (12 PM; **C**) and midactive phases (12 AM; **D**) are shown. Phrenic amplitude as a percent change from baseline (% pLTF) at 90 min post 3×5 mAIH was not different between the midrest and midactive phases (n=7 each group; p = 0.430; two-way RM ANOVA). Bars show mean ± SEM. **+** p < 0.001, significant differences *vs* baseline. Hx, hypoxia; FiO_2_, fraction of inspired oxygen; mAIH, moderate acute intermittent hypoxia; pLTF, phrenic long-term facilitation.

We originally hypothesized that serotonin-dependent mAIH-induced pLTF would be lower in the active *versus* rest phase due to elevated background adenosine levels. However, despite measured increases in spinal adenosine during the active phase, pLTF magnitude was similar during midrest (56.1 ± 5.2% pLTF; n = 7) *versus* midactive phase (50.5 ± 4.4% pLTF; n = 7) phase (F_1,27_ = 0.667, p = 0.430, two-way RM ANOVA; **Figure 3C-E**). Integrated phrenic nerve burst amplitude 90 min post-3×5 mAIH was significantly elevated *versus* baseline activity, indicating significant pLTF (F_1,12_ = 245.994, p < 0.001, two-way RM ANOVA), during midrest and midactive phases (both p < 0.001; Tukey *post-hoc*). Since pLTF magnitude was unchanged by time-of-day despite elevated basal adenosine levels in the active phase, we wondered if the combined time-of-day and 5 minute hypoxic episodes effects were sufficient to flip pLTF from serotonin- to adenosine-dominance.

### 5 min moderate hypoxic episodes in the active phase flips pLTF from serotonin to adenosine dominance

To address the possibility that there is a shift in mechanism from Q (serotonin) to S (adenosine) pathway dominance with time-of-day, we hypothesized that cervical spinal A_2A_ receptor inhibition would impair pLTF in the midactive phase, even though it is known to augment mAIH-induced pLTF in the rest phase ^23^. The A_2A_ receptor antagonist, MSX-3 (130 ng/kg, 10 μM, 12 μL), was delivered intrathecally at C_4_ to localize inhibition near the phrenic motor nucleus ^14,23^ 12 minutes prior to 3×5 min mAIH to determine if increased adenosine during the active phase (plus hypoxia evoked adenosine release) is sufficient to flip mAIH-induced pLTF from a serotonin- to an adenosine-driven mechanism (**Figure 4A**).

**Figure 4.**
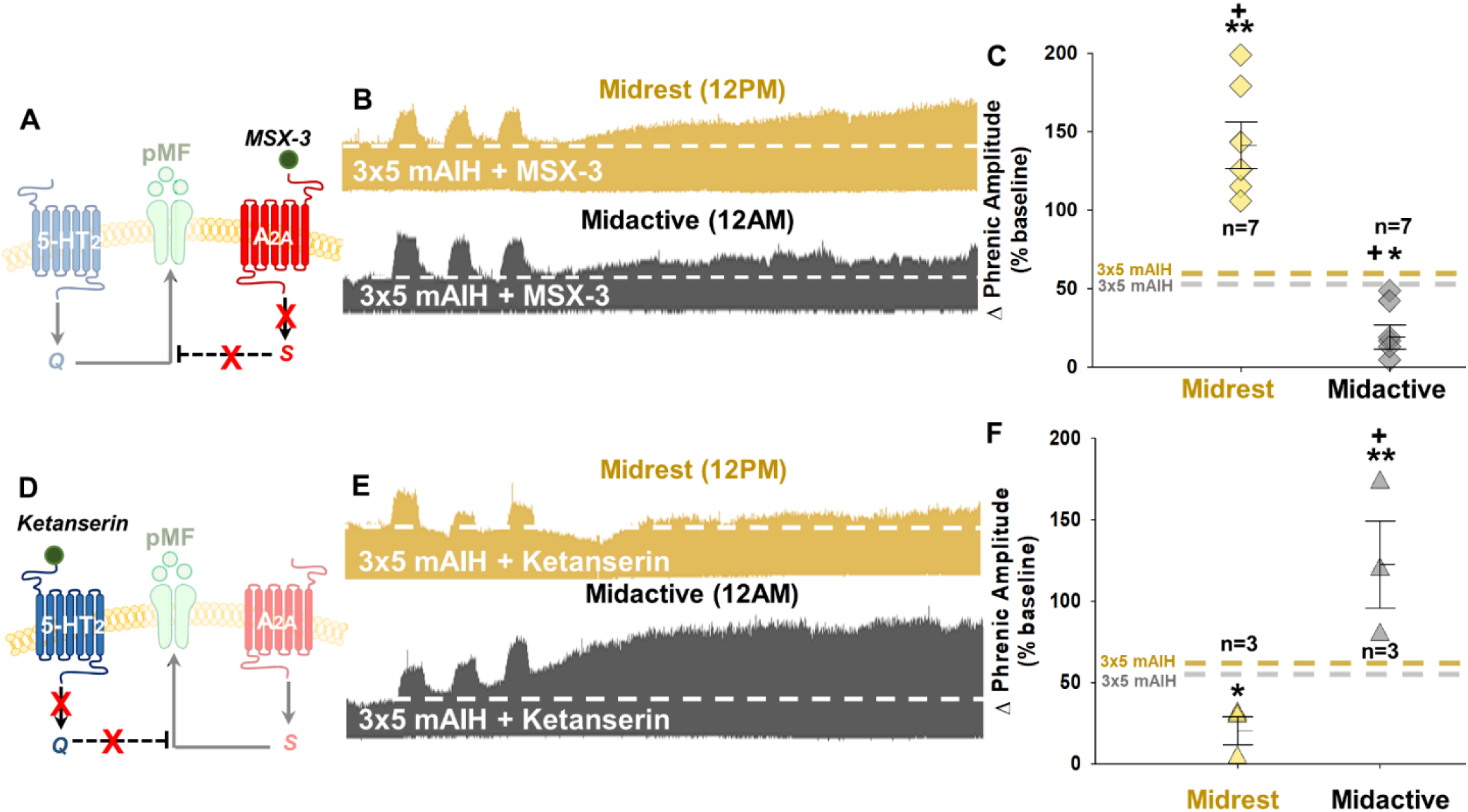
3×5 mAIH induced pLTF flips from serotonin-dominant in the midrest to adenosine-dominant in the midactive phase. The adenosine 2A receptor (A_2A_) inhibitor MSX-3 was administered intrathecally at C_4_ 15 min prior to 3×5 mAIH to minimize the adenosine constraint of serotonin-dominant plasticity during midrest (**A**). Representative compressed neurogram of integrated phrenic nerve activity after intrathecal MSX-3 during baseline, 3×5 mAIH (Hx1-3; PaO_2_ = 40-55 mmHg), and 90 min post-mAIH (**B**) at midrest (12 PM; *top*) and midactive (12 AM; *bottom*) phase. During midrest, A_2A_ inhibition prior to 3×5 mAIH increases pLTF (threeway RM ANOVA, Tukey *post-hoc; vs* 3×5 mAIH alone, p < 0.001); in contrast, pLTF is significantly reduced by the same treatment in the midactive phase (*vs* 3×5 mAIH alone, p = 0.007; *vs* 3×5 mAIH + MSX-3 at midrest, p < 0.001; **C**). To verify pLTF mechanisms shift between rest and active phases, the 5-HT_2A_ receptor inhibitor, ketanserin, was given intrathecally 15 min before 3×5 mAIH (**D**). Compressed neurograms of integrated phrenic nerve activity after intrathecal ketanserin delivery during baseline, 3×5 mAIH protocol and 90 min post-mAIH (**E**) are shown in the midrest (12 PM; *top*) and midactive phases (12 AM; *bottom*). During midrest, 5-HT_2A_ receptor inhibition before 3×5 mAIH suppresses pLTF (3-way RM ANOVA, Tukey *post-hoc; vs* 3×5 mAIH alone, p < 0.022); the same treatment significantly enhances pLTF in the midactive phase (*vs* 3×5 mAIH alone, p < 0.001; *vs* 3×5 mAIH + ketanserin, p < 0.001; **F**). Bars indicate mean ± 1 SEM; significant differences: *p < 0.030 *vs* 3×5 mAIH at same time of day; **p < 0.030 *vs* 3×5 mAIH at same time of day and *vs* midactive 3×5 mAIH + MSX-3 (**C**) *or* midrest 3×5 mAIH + ketanserin (**F**); **+** p < 0.001 *vs* baseline.

Although baseline phrenic nerve activity was unaffected by drug administration, 3×5 mAIH elicited significant pLTF, and time-of-day and MSX-3 significantly altered the magnitude of pLTF in the overall ANOVA (F_1,24_ = 52.27, p < 0.001, three-way RM ANOVA; **Figures 4B** and **C**). MSX-3 pretreatment signicantly enhanced 3×5 mAIH-induced pLTF during **<u>midrest</u>** (141.4 ± 14.8% pLTF; n = 7) *versus* vehicle pretreatment (p < 0.001, Tukey *post-hoc*) and baseline (p < 0.001, Tukey *post-hoc*), confirming prior reports ^16,23^. In contrast, when 3×5 mAIH plus MSX-3 were delivered in the **<u>midactive</u>** phase, pLTF was significantly reduced (19.1 ± 7.6% pLTF; n = 7) *versus* 3×5 mAIH alone (p = 0.007, Tukey *post-hoc*; **Figures 4B** and **C**). These reverse effects of cervical spinal A_2A_ receptor inhibition on 3×5 mAIH-induced pLTF in the midrest versus midactive phases strongly suggest a flip from a serotonin-dominant, adenosine constrained (midrest) to an adenosine-dominant, serotonin constrained mechanism. This mechanistic flip likely results from combined background (basal) adenosine levels and hypoxia-induced adenosine release during the midactive phase.

To further verify this hypothesis, the serotonin 2A/2C (5-HT_2A/2C_) receptor antagonist, ketanserin tartrate (500 μM, 18 μL), was delivered intrathecally at C_4_ to block serotonin-dependent Q pathway activation (**Figure 4D**). There were significant effects of 3×5 mAIH, time-of-day and ketanserin on pLTF (F_1,16_ = 29.187, p < 0.001, three-way RM ANOVA). During midrest, ketanserin pretreatment (n = 3) reduced pLTF to 20.2 ± 8.6% (p = 0.022, Tukey *post-hoc*; **Figures 4E** and **4F**), and it was not significantly different from baseline (p = 0.680, Tukey *post-hoc*), confirming previous reports ^15^. In striking contrast, ketanserin pretreatment during the midactive phase enhanced pLTF (123 ± 27%; n = 3) *versus* 3×5 mAIH alone (p < 0.001, Tukey *post-hoc*; **Figures 4E** & **4F**). Thus, although 3×5 mAIH induces the *same* pLTF magnitude in each phase of the day, it does so by an *entirely different mechanism*, shifting from serotonin-dominant, adenosine constrained in the midrest to adenosine dominant, serotonin constrained in the midactive phase.

### Shorter hypoxic episodes minimize evoked adenosine release in phrenic motor nucleus

Hypoxic episodes elicit rapid carotid chemoreceptor activation and subsequent brainstem serotonergic neuron activation ^3,11,47^. In contrast to rapid carotid chemoreceptor and neural network activation, hypoxia spreads to the spinal cord tissues more slowly, where it triggers glial ATP release with subsequent conversion to adenosine. Thus, hypoxia-evoked adenosine release is slower than neural network activation which triggers serotonin release and the Q pathway to phrenic motor facilitation. Based on these time course differences, shorter hypoxic episodes may emphasize serotonergic *versus* adenosinergic effects on phrenic motor plasticity. Five minute hypoxic episodes give adequate time to reach a nadir in spinal tissue hypoxia ^48^; however, it is possible that parsing the same cumulative duration of hypoxia (15 minutes) into shorter and more numerous episodes (1 min) may minimize hypoxia-evoked adenosine release, shifting the serotonin/adenosine balance in favor of serotonergic mechanisms regardless of time of day.

To test this hypothesis, spinal tissue PO_2_ was measured using a 50 μM optical sensor (Unisense; Aarhus, Denmark) positioned 1mm lateral to midline between C_3_ and C_4_, approximately 1.5 mm below the surface of the spinal cord (**Figure 5A;** *see* ^48^). We then presented 1 *versus* 5 min moderate hypoxic episodes, with the inspired O_2_ adjusted to achieve the same spinal tissue PaO_2_ nadir within episodes. Hypoxia was delivered as: 1) 5 min of 13% inspired O_2_ (PaO_2_ = 46.6 ± 3.9 mmHg; **Figure 5B**); or 2) 1 min at 9% inspired O_2_ (PaO_2_ = 46.2 ± 1.3 mmHg; **Figure 5C;** n = 4 per group). In these rats, no differences in baseline tissue PO_2_ were observed (t_6_ = 0.12, p = 0.912; unpaired *t*-test; **Figure 5D**). During 5 *versus* 1 min episodes, spinal PtO_2_ achieved similar levels during 5 min episodes despite differences in inspired oxygen concentration: PtO_2_ = 13.8 ± 2.8 mmHg and 12.5 ± 1.2 mmHg for 5 and 1 min hypoxic episodes, respectively (t_6_ = 0.42, p = 0.689; unpaired *t*-test; **Figure 5E**). Tissue reoxygenation was similar after both 5 and 1 min episodes (t_6_ = −0.44, p = 0.678; unpaired *t*-test; **Figure 5F**).

**Figure 5.**
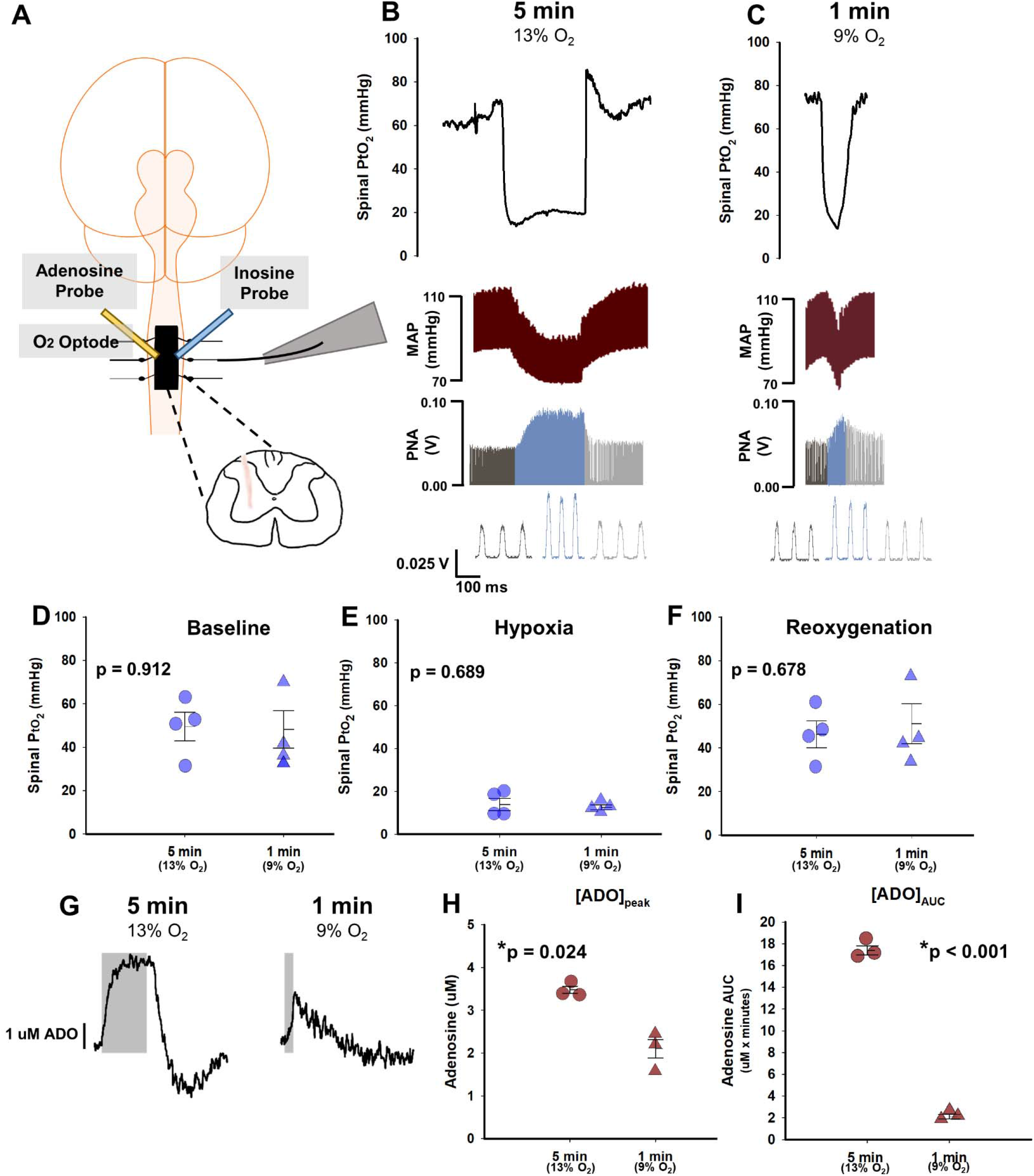
One minute episodes of 9% O_2_ reach the same level of tissue hypoxia as 5 minute, 13% O_2_ episodes, yet these shorter episodes evoke less adenosine accumulation. In separate experiments, either an oxygen optode or adenosine/inosine probes were placed between ventral C_3_/C_4_ to measure changes in tissue oxygen and adenosine accumulation, respectively (**A**). During 5 (13% O_2_; **B**) *versus* 1 min episodes (9% O_2_; **C**), representative traces illustrate similar nadirs in spinal tissue oxygen pressure (PtO_2_), as well as peaks in mean arterial pressure (MAP) and phrenic nerve amplitude (PNA). There was no difference in mean spinal PtO_2_ between 5 *versus* 1 min hypoxic episodes during baseline (**D**), hypoxia (**E**) or reoxygenation (**F**) (n = 4 per group; unpaired *t*-test). Despite similar reductions in arterial PO_2_ and PtO_2_ during 5 min versus 1 min episodes, greater adenosine accumulation (μM) was observed in 5 min episodes (**G**; n = 3 per group; unpaired *t*-test), either when expressed as peak adenosine level ([ADO]_peak_; **H**) or total area under the curve ([ADO]_AUC_; **I**). Bars are means ± 1 SEM; *p < 0.030.

Evoked adenosine release was also measured in the ventral cervical spinal cord near the phrenic motor nucleus during 5 and 1 minute hypoxic episodes *via* adenosine and inosine microbiosensors (Zimmer-Peacock, UK; n = 3 per group; **Figure 5G**). With 5 minute moderate hypoxic episodes, significantly higher peak adenosine levels were observed *versus* 1 min episodes (5 min: 3.5 ± 0.1 μM; 1 min: 2.1 ± 0.2 μM; t_4_ = 6.04, p = 0.004; unpaired *t*-test; **Figure 5H**). When expressed as area under the curve to reflect total adenosine accumulation, significantly more adenosine accumulated with 5 *versus* 1 min hypoxic episodes (5 min: 17.4 ± 0.4 μM; 1 min: 2.1 ± 0.21 μM; t_4_ = 33.10, p < 0.001; unpaired *t*-test; **Figures 5I**). Thus, shorter hypoxic episodes trigger less adenosine release near phrenic motor neurons despite similar levels of spinal tissue hypoxia.

Collectively, these data support the idea that adenosine accumulation during 5 min hypoxic episodes, when superimposed on elevated basal adenosine levels in the active phase, provide sufficient adenosine to ‘push’ 3×5 mAIH-induced pLTF from a serotonin-dominant to adenosine-dominant plasticity. From another perspective, these findings strongly suggest that one way to minimize the undermining influences of hypoxia-evoked adenosine release is to shorten hypoxic episodes, even with the same tissue oxygen levels. Thus, we tested the hypothesis that 15, 1 min moderate hypoxic episodes elicit greater pLTF during the rest phase versus the standard 3, 5 min mAIH protocol, despite the same cumulative duration of hypoxia (15 minutes).

### Shorter hypoxic episodes amplify pLTF during midrest but attenuate pLTF in the midactive phase

To minimize adenosine accumulation during mAIH, hypoxic episode duration was shortened to 15, 1 min hypoxic episodes with 1 min intervals of baseline conditions in the midrest and midactive phases (**Figure 6A**). The impact of 15×1 mAIH on pLTF is time of day dependent (F_1,27_ = 54.814, p < 0.001, two-way RM ANOVA; **Figures 6B** and **6C**). Although phrenic amplitude 90 min post-mAIH was significantly elevated versus baseline in midrest and midactive phases (p < 0.001 and p = 0.013, respectively; Tukey *post-hoc*), the 15×1 mAIH protocol elicited enhanced pLTF during midrest (138.6 ± 14.0% pLTF; n = 7), reaching levels comparable to 3×5 mAIH after spinal A_2A_ receptor inhibition (**Figure 4A**). In striking contrest, this same mAIH protocol applied during the midactive phase elicits diminished pLTF (30.4 ± 4.1% pLTF; n = 7); during the midactive phase, 15×1 mAIH-induced pLTF was significantly less than midrest (p < 0.001; Tukey *post-hoc*), and significantly less than pLTF elicited by 3×5 mAIH during the midactive phase (p = 0.021). Thus, although 15×1 mAIH amplifies pLTF during midrest, when background adenosine levels are low, pLTF is constrained during the midactive phase, when background adenosine is elevated. Phrenic amplitude 90 min post-mAIH was significantly elevated versus baseline in both midrest and midactive phases (p < 0.001 and p = 0.013, respectively; Tukey *post-hoc*); however, in striking contrast to the lack of change in 3×5 mAIH-induced pLTF during midactive *versus* midrest phases, 15×1 mAIH exhibits a striking time-of-day effect. We suspected that this time-of-day effect can be accounted for by differential adenosine constraints of serotonin-dominant pLTF in the rest versus active phases of the day.

**Figure 6.**
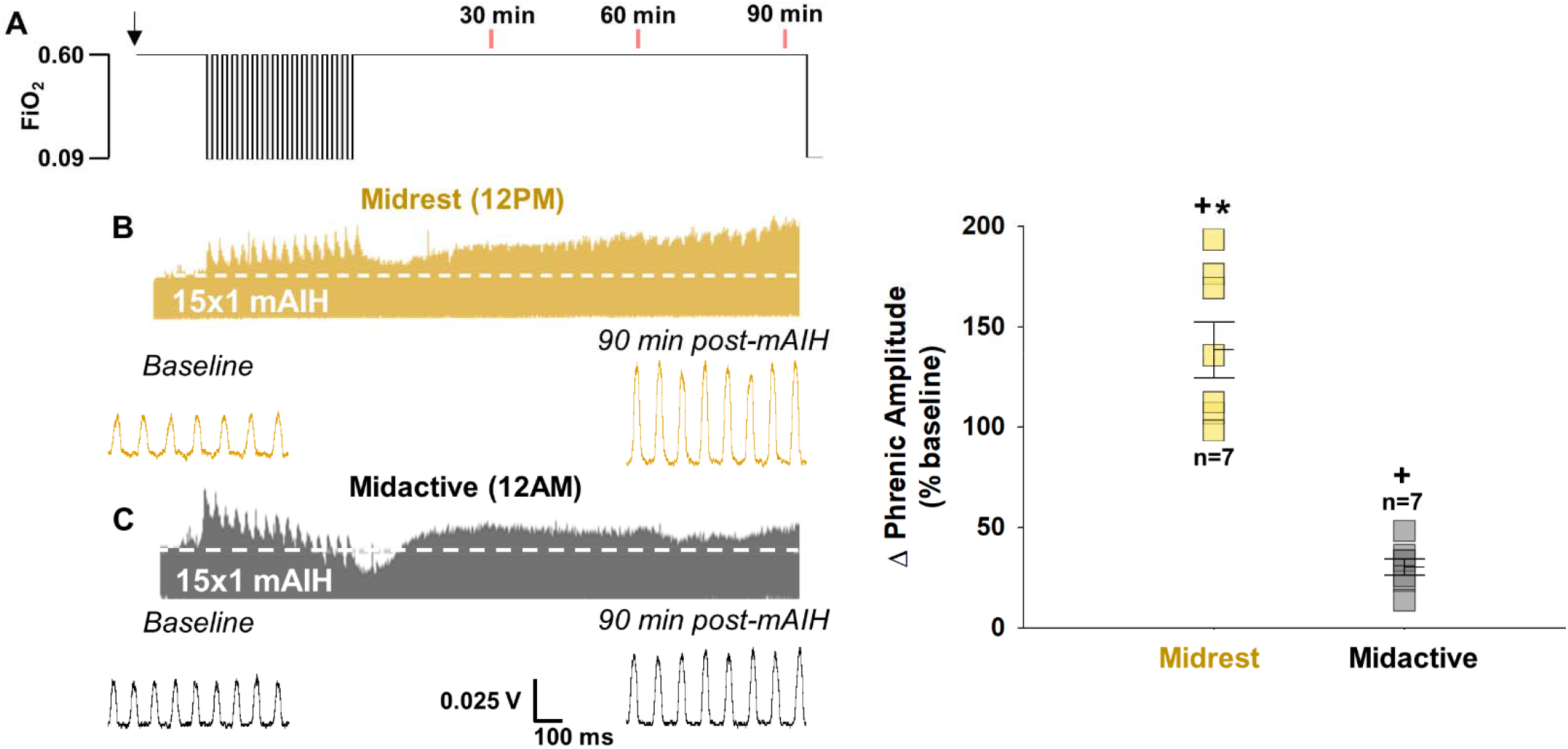
With 15×1 mAIH, pLTF is enhanced during midrest (noon) but suppressed during midactive phase (midnight). With the 15×1 mAIH protocol (**A**), compressed integrated phrenic responses are shown during baseline, 15×1 mAIH, and 90 min post-mAIH at midrest (12 PM; **B**) and midactive (12 AM; **C**) phase. Phrenic burst amplitude, represented as percent change from baseline at 90 min post 15×1 mAIH, is substantially different during midrest vs. midactive phases (n=7 per group; three-way RM ANOVA, p < 0.001). Bars show means ± 1 SEM. Significant differences for: *p < 0.001, *vs* midactive 15×1 mAIH; **+**p < 0.020, *vs* baseline. FiO_2_, fraction of inspired oxygen; mAIH, moderate acute intermittent hypoxia.

To verify that this striking difference in 15×1 mAIH-induced pLTF with time of day was due to differences in background adenosine levels, MSX-3 was delivered intrathecally at C_4_ to block cervical spinal A_2A_ receptors prior to administering 15×1 mAIH (n = 7 per group). There was a significant interaction between time of day and MSX-3 pretreatment on pLTF (F_5,28_ = 12.05, p < 0.001; three-way mixed effects ANOVA; **Figures 7A** and **7B**). Specifically, MSX-3 had no significant effect on 15×1 mAIH-induced pLTF during midrest (121.5 ± 10.7% pLTF; p = 0.931; Tukey *post-hoc*), confirming the robust midrest pLTF (versus 3×5 mAIH) results from diminished hypoxia-evoked adenosine accumulation at a time with low background adenosine levels. In striking contrast, A_2A_ receptor inhibition prior to 15×1 mAIH during the midactive phase greatly enhanced pLTF (p < 0.001 *vs* without MSX-3; Tukey *post-hoc*), reaching levels consistent with those observed during midrest (163.7 ± 26.7% pLTF; p = 0.591; **Figures 7B** and **7C**). To assure proper receptor targeting by MSX3, a second selective A_2A_ receptor inhibitor, istradefylline, was used to verify MSX-3 effects (n = 3); istradefylline pretreatment during the midactive phase also increased 15×1 mAIH-induced pLTF *versus* 15×1 mAIH alone (p < 0.001; Tukey *post-hoc*), consistent with MSX-3 (193.1 ± 56.6% pLTF; p = 0.882, Tukey *post-hoc*; **Figure 7B** and **7C**).

**Figure 7.**
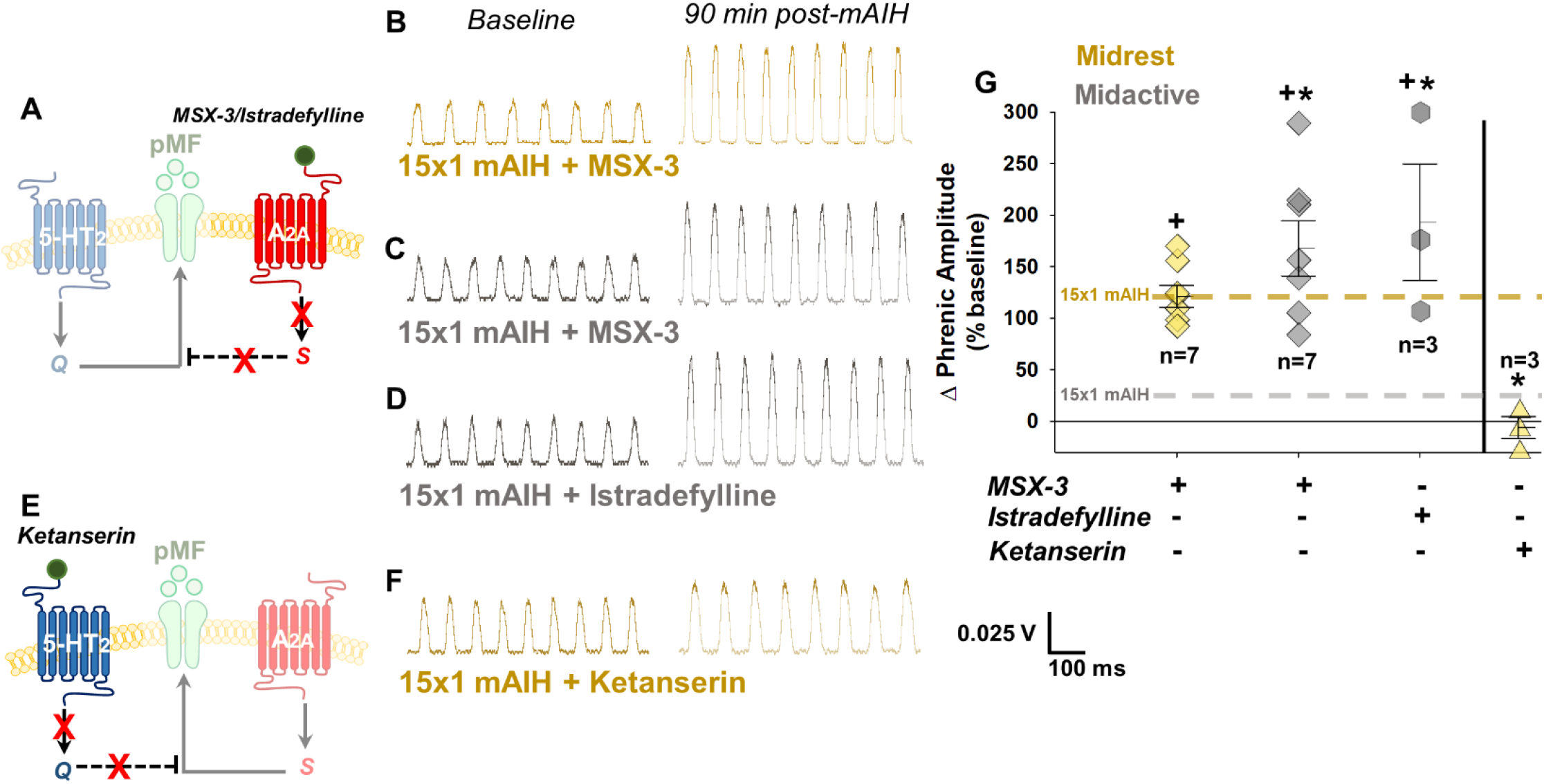
Following 15×1 mAIH, pLTF is serotonin-driven during both midrest and midactive phases. Adenosine 2A receptor (A_2A_) inhibition with MSX-3 (intrathecal at C_4_ 15 min prior to 15×1 mAIH) was used to minimize any adenosinergic constraint on serotonin-driven plasticity (**A**). Compressed integrated phrenic neurograms after intrathecal MSX-3 are shown during baseline and 90 min post-mAIH for midrest (12 PM; **B**) and midactive phase (12 AM; **C**). Compressed i integrated phrenic neurograms are shown after intrathecally delivered istradefylline, another selective A_2A_ receptor inhibitor, during baseline and 90 min post-mAIH during the midactive phase (**D**). Representative neurograms of integrated phrenic nerve activity after intrathecal ketanserin during baseline and 90 min post-15×1 mAIH at midrest confirm serotonin-dependent pLTF during midrest and midactive phases (**F**). During midrest, A_2A_ receptor inhibition prior to 15×1 mAIH (n = 7) had no effect on pLTF (three-way mixed effects ANOVA, Tukey *post-hoc; vs* 15×1 mAIH alone, p = 0.931; **G**), although pLTF was restored in the midactive phase (n = 7) to levels observed during midrest (*vs* 15×1 mAIH alone, p < 0.001; *vs* 15×1 mAIH + MSX-3 at midrest, p = 0.123); A_2A_ receptor specificity was confirmed with istradefylline (n = 3; midactive 15×1 mAIH + MSX-3 *vs* istradefylline, p = 0.882). Further, pLTF during midrest 15×1 mAIH was significantly reduced by ketanserin *versus* midrest 15×1 mAIH alone (n = 3; p < 0.001). Bars show means ± 1 SEM; Significant differences: *p < 0.001 *vs* respective midrest or midactive 15×1 mAIH; **+**p < 0.001 *vs* baseline.

Finally, intrathecal ketanserin was administered to block cervical spinal 5-HT_2A/C_ receptors prior to 15×1 mAIH during midrest (n = 3) to confirm that pLTF was driven by the serotonin-dependent Q pathway. As expected, ketanserin pretreatment during midrest blocked pLTF (−7.9 ± 10.8% pLTF; p = 0.692 *vs* baseline; Tukey *post-hoc*), and was significantly lower than 15×1 mAIH alone (p < 0.001; Tukey *post-hoc*; **Figures 7D** and **7E**). Thus, mAIH consisting of shorter hypoxic episodes drives serotonin-dominant plasticity during both rest and active phases, although elevated adenosine during the active phase undermines serotonin-dominant plasticity.

### Low (basal and evoked) adenosine levels allow pLTF to ‘build’ within the mAIH protocol

Over years of study using the 3×5 mAIH, pLTF has been reported to build during the successive intervals (Mitchell et al., 2001; Bach and Mitchell, 1996). However, with the shorter hypoxic episodes of the 15×1 mAIH protocol, stepwise increases in phrenic burst amplitude were observed during the inter-hypoxic intervals (pLTF ‘building’). There is a strong correlation between the change in integrated phrenic nerve burst amplitude during inter-hypoxic intervals and pLTF at 90 min post-mAIH.

The serotonin-dependent, Q pathway to phrenic motor facilitation requires ERK activation, new BDNF protein synthesis and TrkB activation within phrenic motor neurons ^18,19,49–51^. Since 3×5 mAIH-induced pLTF requires rapid ERK MAPK activation ^49,52^, and new BDNF protein synthesis requires 15-20 minutes poststimulus ^53^, pLTF ‘building’ likely reflects an early processes in pLTF, such as ERK MAPK activation. Changes in integrated phrenic nerve burst amplitude were averaged during successive inter-hypoxic intervals following the first hypoxic episode (**Figure 8** and **S3-S4**). Consistent with earlier reports ^11,54–56^, increases in phrenic nerve burst amplitude were not observed during the inter-hypoxic intervals of the 3×5 mAIH protocol (**Figure S3**), regardless of time of day (n = 7 each group; F_1,6_ = 1.341, p = 0.291; one-way RM ANOVA) or drug pretreatment (n = 3-7 each group; midrest: F_2,33_ = 1.485, p = 0.260; midactive: F_2,33_ = 1.592, p = 0.238; threeway RM ANOVA). Further, there was no relationship between activity during these intervals and pLTF 90 min post-mAIH (**Figure S4**; p > 0.050).

**Figure 8.**
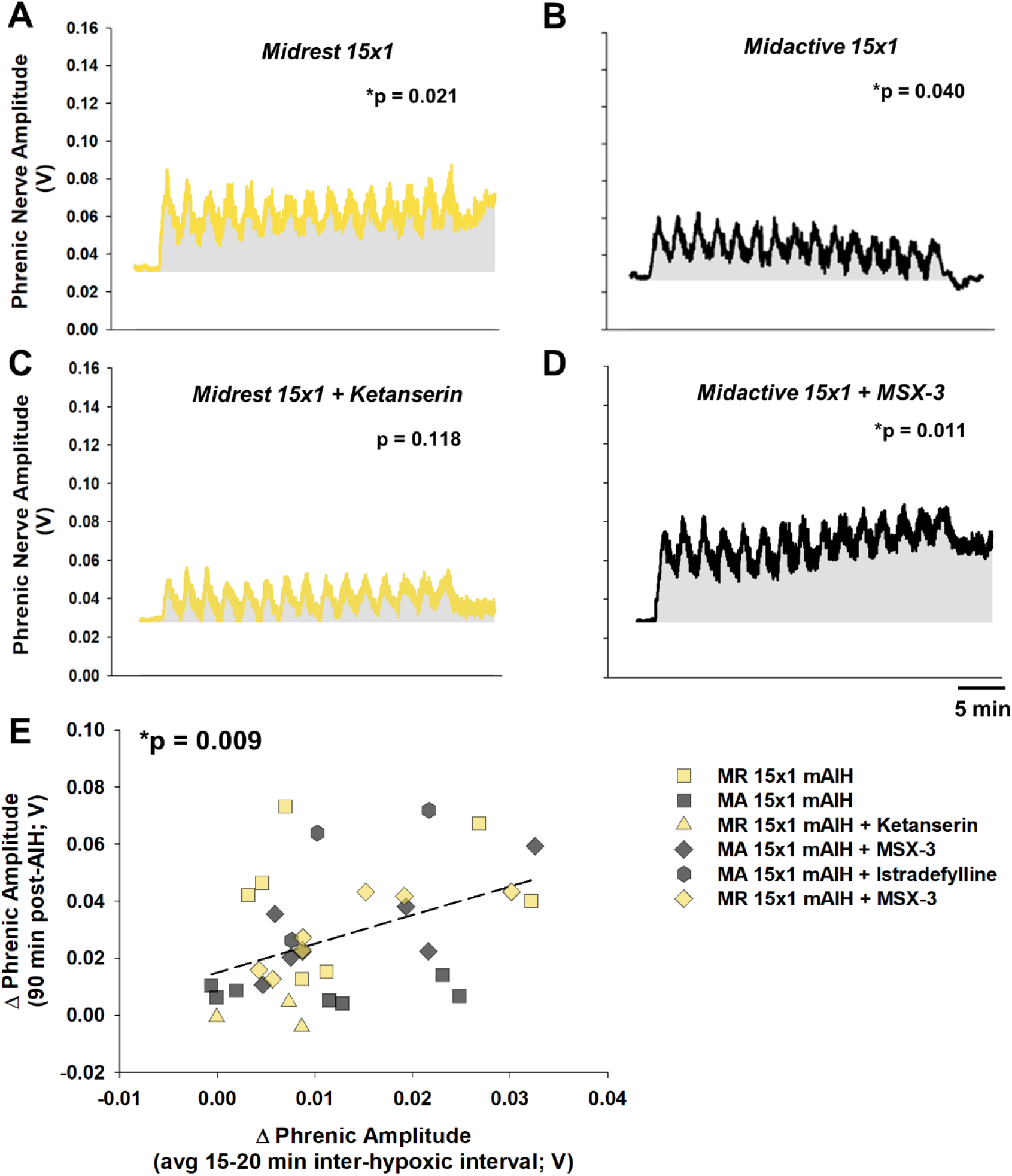
pLTF ‘building’ in 15×1 mAIH inter-hypoxic intervals requires serotonin receptor activation and is impaired by spinal adenosine. Further, pLTF ‘building’ correlates with pLTF 90 min post-AIH. The 15×1 mAIH protocol exhibits pLTF ‘building’ during midrest and midactive phases, although less prominent during the mid-active phase; average traces of 15×1 mAIH during midrest (n = 7; **A**) and midactive (n = 7; **B**) phases. After pretreating with ketanserin, pLTF ‘building’ is blocked during midrest (n = 3; **C**). MSX-3 reveals pLTF ‘building’ during the midactive phase (n=7; **D**). Gray area represents area-under-curve as a change from baseline. *p < 0.050, significant difference in phrenic nerve activity from baseline (one-way RM ANOVA). In **E**, shorter hypoxic durations and serotonin and adenosine receptor inhibition modify correlation between pLTF ‘building’ during 15×1 mAIH and pLTF 90 min post-AIH (regression analysis; p = 0.009).

In striking contrast, when the mAIH consisted of 15, 1 min hypoxic episodes, stepwise increases in phrenic nerve burst activity were observed in the inter-hypoxic intervals. Phrenic nerve amplitude significantly increased from baseline in successive intervals during the 15×1 mAIH protocol during midrest (n = 7; F_1,6_ = 9.620, p = 0.021; one-way RM ANOVA), when basal spinal adenosine is low (**Figure 8A**). In comparison, more modest changes were observed in the midactive phase, when basal spinal adenosine levels are elevated; the rise during successive intervals consisted of initial facilitation with successive decreases as the mAIH protocol progressed (**Figure 8B**; n = 7; F_1,6_ = 6.863, p = 0.040; one-way RM ANOVA). During the rest phase, ketanserin blocked pLTF ‘building’ during the inter-hypoxic intervals (**Figure 8C**; n = 3; F_1,2_ = 3.881, p = 0.188; one-way RM ANOVA).

Progressive increases in phrenic nerve activity during the interhypoxic intervals of the 15×1 mAIH protocol are consistent with reports that phrenic nerve activity ramps up in successive intervals between episodes of carotid sinus nerve electrical stimulation in cats (where there is no hypoxia), and that the stepwise increases in phrenic nerve activity between episodes of carotid sinus nerve stimulation are serotonin-dependent ^7,9,12^. In this case, episodic electrical activation of the carotid sinus nerve (i.e. brainstem neural network activation) drives serotonin-dependent phrenic motor plasticity in the absence of hypoxia-induced adenosine accumulation. In the midactive phase, we suggest that basal spinal adenosine levels are sufficient to prevent pLTF “building” during 15×1 mAIH. Consistent with this hypothesis, significant inter-episode increases in phrenic nerve amplitude were observed when A_2A_ receptors were blocked in the midactive phase (**Figure 8D**; MSX-3: n = 7; F_1,6_ = 13.269, p = 0.011; one-way RM ANOVA). In this case, changes in phrenic nerve burst amplitude during the inter-hypoxic intervals of the 15×1 mAIH protocol correlate with pLTF at 90 min post-mAIH (**Figure 8E**; p = 0.009). Thus, by suppressing the adenosine constraint during mAIH via shorter hypoxic episodes in the midrest phase (when basal spinal adenosine is low) or cervical spinal A_2A_ receptor inhibition (when basal spinal adenosine is high), we revealed a profound increase in serotonin-driven phrenic motor plasticity. These data provide unique insights concerning how protocol specificity plays an important role in the early signaling events associated with mAIH-induced pLTF.

## DISCUSSION

We demonstrate that adenosine is a powerful regulator of mAIH-induced phrenic motor plasticity across the daily rest/active cycle. In combination, basal and hypoxia-evoked adenosine regulate both the magnitude and mechanism of plasticity. Since AIH-induced pLTF is an important model of respiratory motor plasticity in a critical neural system, knowledge of factors regulating pLTF is of considerable importance to both future experimental design and our understanding of its biological/physiological significance. Further, since increased understanding of pLTF mechanisms is already guiding efforts to harness AIH as a therapeutic modality to restore breathing and other movements in people with chronic spinal cord injury and ALS ^3,57–59^, the findings reported here represent a “game changer” since they make clear that both time of day and protocol are factors that must be considered before we will ever be able to optimize this promising therapeutic modality.

We propose that daily rest/active cycles shift both the magnitude and mechanism of pLTF in a protocol-specific manner based on a multi-disciplinary approach. Methods used here include: 1) direct measurements of both basal and evoked adenosine and serotonin levels in the cervical spinl cord (and their respective receptors); 2) direct mAIH-induced pLTF measurements in rats; and 3) pharmacological manipulations demonstrating that the mAIH protocol *and* time-of-day exert powerful influences on both the magnitude and even dominant mechanism driving pLTF. Collectively, these approaches all point to a new conceptual framework concerning the dynamical interplay between chemoreceptor activated neural networks (serotonin) *versus* local tissue oxygen effects of ATP/adenosine release in AIH-induced phrenic motor plasticity. Since new framework includes previously unrecognized (and powerful) time-of-day and mAIH protocol effects, the results of this study constitute a major advance in our understanding of spinal motor plasticity. Going forward, it is essential to consider protocol specific time-of-day effects in the design of experiments, as well as clinical trials attempting to harness AIH as a therapeutic modality to restore breathing and other movements (such as limb function) in people living with chronic spinal cord injury, ALS and a range of other neuromuscular disorders.

### Impact of adenosine on pLTF

Moderate AIH-induced pLTF is an important model of spinal motor plasticity with considerable biological and translational significance. During the past two decades, considerable advances have been made in our mechanistic understanding of mAIH-induced plasticity in the phrenic motor system, a motor system once thought of as fixed and immutable. Despite such progress, virtually no attention has been given to potential time-of-day effects on plasticity, let alone a complete reversal in its underlying mechanisms. Since rodents are nocturnal, studies in rodent models pose a dilemma since humans are diurnal, operating with a reverse circadian cycle. Consequently, even though most rodent and human studies occur during the day, they are in opposite activity states—the rodent rest phase *versus* the human active phase. Our present findings make clear that this oversight could have major implications for our understanding and application of AIH-induced motor plasticity.

During the rest phase, basal spinal adenosine levels are low (**Figure 2**), and peak extracellular adenosine levels are reliant on the duration and severity of hypoxic episodes within the AIH protocol (i.e., 3×5 *versus* 15×1 min episodes). Since shorter hypoxic episodes minimize evoked adenosine release (**Figure 5**), the adenosine constraint on serotonin-dominant plasticity is minimized by using 15×1 *versus* 3×5 mAIH. After pretreatment with an A_2A_ receptor inhibitor during the rest phase, 3×5 mAIH-induced pLTF is similar in magnitude to that elicited by 15×1 mAIH withough pretreatment (**Figures 4** and **7**). In contrast, elevated background adenosine levels suppress pLTF when using 15×1 mAIH during the active phase (**Figure 6**). Even more striking, the “standard” 3×5 mAIH protocol triggers greater hypoxia-evoked adenosine release that constrains pLTF during midrest. However, this same mAIH protocol combines with elevated background adenosine levels during the active phase, reaching adenosine levels sufficient to flip from serotonin-driven (midrest) to adenosine-driven plasticity in the midactive phase (**Figure 3**). Thus, depending on time of day and mAIH protocol, adenosine constrains or dominates phrenic motor plasticity, a finding unique among studies of neuroplasticity.

Adenosine links brain activity state with the sleep-wake cycle and neuroplasticity ^31^. With spinal respiratory neuroplasticity, the impact of adenosine across the daily rest/active cycle was previously unknown. We now demonstrate a fundamental link between daily *spinal* adenosine fluctuations and phrenic motor plasticity, and that its impact is modified by hypoxia evoked-adenosine release and accumulation. Adenosine influences pLTF via A_2A_ receptor activation, initiating intracellular signaling cascades that inhibit the serotonin-dependent Q pathway, or drive a distinct mechanism of plasticity, the S pathway to phrenic motor facilitation ^16,18,54,60^.

Although hypoxia-evoked adenosine release from local glia and its accumulation has been proposed to depend on the severity and duration of hypoxic episodes based on indirect evidence ^16,22^, we now directly demonstrate greater adenosine accumulation during longer (5 min) versus shorter (1 min) hypoxic episodes, despite similar levels of tissue hypoxia in these experiments (**Figure 5**). Evoked adenosine release occurs on shifting baseline adenosine levels across time-of-day. Although it is well known that brain extracellular adenosine levels increase during prolonged wakefulness, and decrease during sleep/rest ^26,41,61^, we directly demonstrate similar time-dependent shifts in the ventral cervical spinal cord near the phrenic motor nucleus (**Figure 2**). These findings make clear that it is essential to consider both the baseline and evoked adenosine levels when evaluating adenosine regulation of AIH-induced phrenic motor plasticity. Together, they have potential to undermine or even replace serotonin-dependent plasticity (**Figure 9**).

**Figure 9.**
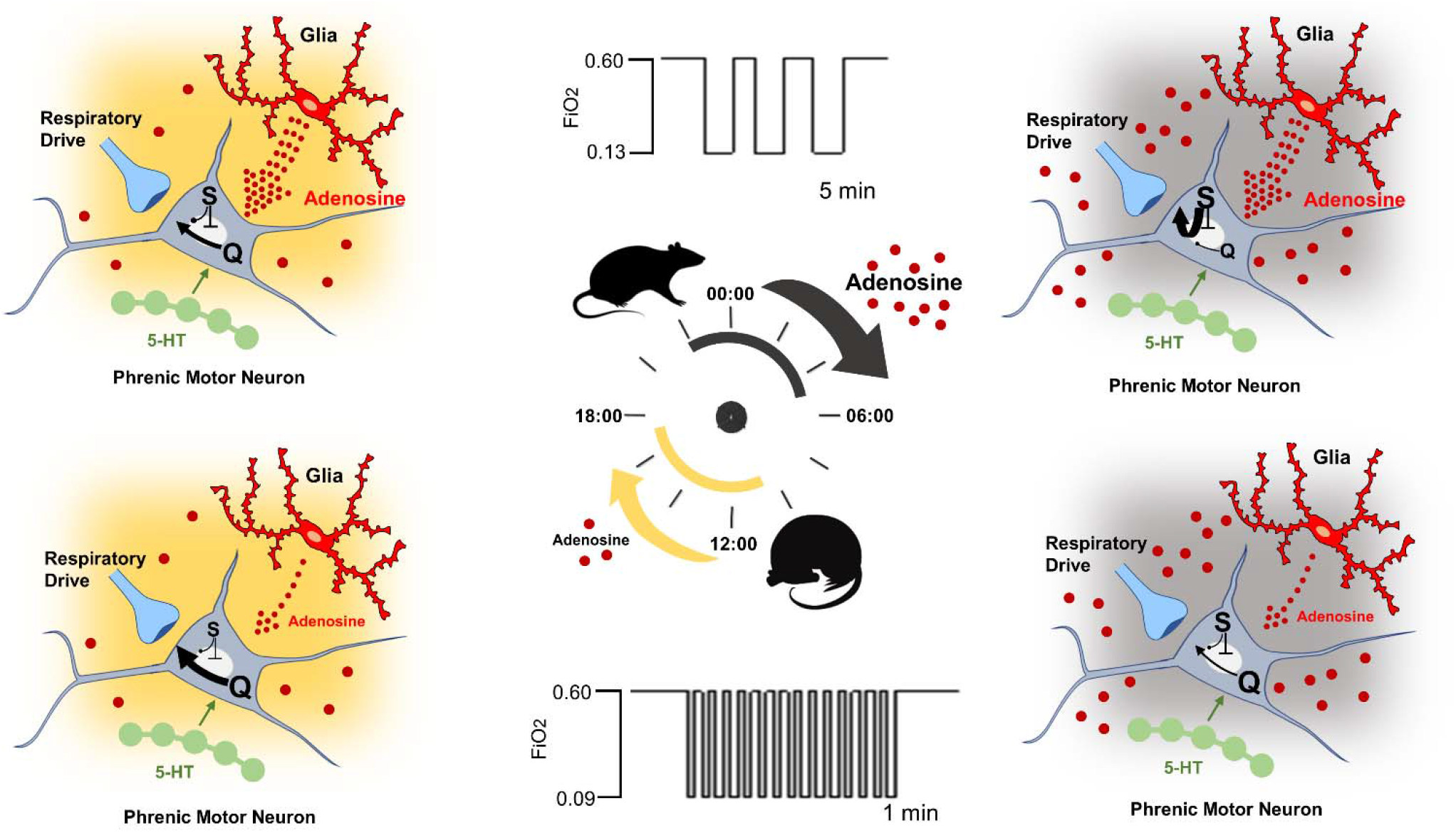
mAIH-induced pLTF is regulated by time-of-day dependent shifts in basal adenosine levels and hypoxic episode duration (i.e., evoked adenosine release). Spinal adenosine (red dots) fluctuates across the daily rest/active cycle, with low levels during the rest phase (yellow) and elevated levels in the active phase (gray background). In the top row, 3, 5 min episodes of moderate hypoxia (3×5 mAIH) evoke greater adenosine release from glia (signified by larger dotted arrows). In the rest phase (*top row, left panel*), 3×5 mAIH induces pLTF through a serotonin-dominant (Q pathway), adenosine constrained mechanism. In the active phase (*top row, right panel*), greater basal adenosine levels near phrenic motor neurons coupled with still high hypoxia-evoked adenosine release cause a mechanistic flip; 3×5 mAIH now induces pLTF through an andenosine-dominant (S pathway), serotonin constrained mechanism. In the bottom row, 15, 1 min episodes of moderate hypoxia (15×1 mAIH) evoke minimal adenosine release (signified by smaller arrow from glia). In the rest phase (*bottom row, left panel*), 15×1 mAIH elicits pLTF through the full-blown serotonin-dependent Q pathway, with minimal adenosine constraint. However, in the active phase (*bottom row, right panel*), 15×1 mAIH still elicits pLTF through the Q pathway, but is now undermined considerably by elevated basal adenosine levels; in this case, 15×1 mAIH does not reach the threshold adenosine level necessary to flip pLTF from the Q to the S pathway, yet the Q pathway is sufficiently undermined to be lower than S-pathway driven 3×5 mAIH-induced pLTF at this time.

Additional triggers to adenosine accumulation in the ventral cervical spinal cord may contribute to the overall ‘adenosine burden’, including neuroinflammation ^62^, aging ^63–65^ or spinal cord injury ^48^. Each of these factors is known to influence mAIH induced phrenic motor plasticity ^66,67^, although the potential role of adenosine in their effects has not yet been explored. As these additional factors are evaluated, time-of-day and the duration of hypoxic episodes in the mAIH protocol will be essential factors to consider.

### The potential to harness serotonin-versus adenosine-dependent plasticity

Although serotonin and adenosine are capable of eliciting phenotypically similar pLTF, it is not yet known if these cellular cascades differentially impact other outcomes, such as axonal growth, synaptogenesis or neuroprotection. Indeed, each plasticity mechanism is driven by molecules associated with neural behaviors such as synaptogenesis ^68^ *versus* axonal growth ^69^. The S pathway requires Akt and mTORC1 signaling ^54,70,71^, a pathway commonly associated with axonal growth/elongation ^69^. Conversely, the Q pathway requires ERK, MAPK and BDNF/TrkB signaling ^49–51,72^, which are more often associated with axon sprouting and synaptogenesis ^68^. The ability to elicit different mechanisms of plasticity based on the time of day and specific AIH protocol applied provides the potential to target these distinct outcomes for more impactful therapeutic outcomes.

### pLTF accumulation during mAIH protocols

Phrenic nerve activity progressively builds during the inter-hypoxic intervals exclusively with the 15×1 mAIH protocol during the rest phase (**Figure 8**). While determining mechanisms of this pLTF ‘building’ was not a focus of this study, our findings are consistent with previous reports using electrical stimulation of the carotid sinus nerve without hypoxia ^9,12^. We suspect that diminished adenosine accumulation characteristic of 15×1 mAIH, in conjunction with low basal adenosine levels during midrest, enables sufficient Q pathway activation to increase phrenic nerve activity. Indeed, pLTF ‘building’ is serotonin dependent with both 15×1 mAIH (**Figure 8**) and episodic carotid sinus nerve stimulation (Millhorn et al., 1980b), suggesting it reflects early stages of Q pathway activation, potentially ERK MAPK activation and/or new BDNF synthesis ^14,49,51,73^. Hypoxia rapidly activates ERK MAPK, initiating BDNF protein synthesis within 15-20 minutes ^53^. Thus, pLTF ‘building’ likely arises from ERK MAPK activity versus BDNF synthesis, which is likely too slow. The strong correlation between pLTF “building’ and pLTF amplitude 90 min post-mAIH regardless of time of day or drug pretreatment is consistent with the idea that these observations reflect the same cascade/mechanism.

Although pLTF ‘building’ is not observed with the 3×5 mAIH protocol, even with A_2A_ receptor inhibition (*see* Supplemental Data), additional factors must override this phenomenon. One potential contributor to this cancellation is greater adenosine accumulation and adenosine 1 receptor activation during hypoxic episodes ^30^.

## LIMITATIONS

Collectively, this body of work represents a unique combination of elements from different fields including *in vivo* neurophysiology, oxygen optodes, adenosine microbiosensors, targeted pharmacologic manipulations of cervical spinal receptors, and complex time-of-day-based physiology. However, since neuromodulators other than adenosine also fluctuate during the sleep/wake cycle, they could provide distinct influences on pLTF; examples of such neuromodulators include corticosterone ^74^, melatonin ^33,75^ and orexin ^76^. Nevertheless, adenosine is likely the primary effector since manipulating A_2A_ receptors had direct effects on pLTF during both midrest and midactive experiments.

All rats used in the present study were male. Female rats are known to exhibit a profound age- and estrus cycle dependent sexual dimorphism in mAIH-induced pLTF ^67,77,78^. It is not known if sex steroid hormones influence pLTF across the daily rest/active cycle. Further, in vivo neurophysiology preparations require anesthesia, which disrupts sleep/wake cycles and, possibly, circadian rhythms ^79,80^. However, urethane is unique among anesthetics since, at least at lower doses, it allows spontaneous shifts between REM-like and nonREM-like EEG states, with corresponding changes in physiological variables typically observed during natural sleep ^81^. From another perspective, we take advantage of the fact that urethane allows long-term stability of physiological state, as required for studies of respiratory neuroplasticity. In conjunction with clear time-of-day shifts in pLTF, molecular data supporting these findings, and care taken to prevent light cues during midactive experiments, the effects reported here are unlikely to be epiphenomena caused by anesthesia. Lastly, unanesthetized preparations have been used to study ventilatory or diaphragm LTF (*vs* pLTF) and, although there is vigilance-state dependence of ventilatory and diaphragm LTF ^82,83^, it is hard to envision how urethane anesthesia could contribute to the time-of-day effects reported here.

## TRANSLATIONAL SIGNIFICANCE

Phrenic LTF is a model of respiratory motor plasticity that has already guided translation in two dimensions: 1) clinical translation to people living with chronic spinal cord injury and ALS (see below); and 2) translation to non-respiratory motor systems (walking, arm/hand function). We are currently engaged in a “translational flywheel,” where rodent experiments drive human studies of therapeutic AIH, which then inform the need for additional rodent studies ^3^. Indeed, multiple funded clinical trials are currently underway, exploring the ability of mAIH to restore breathing, walking and hand/arm function in people living with chronic spinal cord injury and ALS. These clinical trials suffer from the lack of full mechanistic understanding driving optimization of mAIH as a therapeutic modality (*e.g*. optimal mAIH protocol and best time of day to deliver mAIH). Based on the present findings, AIH protocol and time-of-day interactions are important factors to consider. We need to reconsider the current practice of applying therapeutic AIH during the day (the human active phase) in ongoing clinical trials. Alternately, we need to control the influences of elevated adenosine via pretreatment with A2A receptor inhibitors. Consideration of adenosine and time of day in future clinical trials is of major importance.

## CONCLUSION

In summary, we demonstrate shifts in the mechanism of mAIH-induced pLTF between serotonin versus adenosine-dominance depending on time of day and mAIH protocol. These shifts are driven by fluctuations in spinal cord adenosine across the day/night, combined with evoked adenosine release during mAIH. These studies are the first to demonstrate: 1) basal adenosine changes with time of day in spinal tissue containing the phrenic motor system; 2) the powerful role of adenosine in regulating pLTF across the daily rest/active cycle; and 3) the importance of episode duration during mAIH protocols in a manner that depends on time of day. Collectively these findings advance our understanding of mechanisms regulating AIH-induced motor plasticity. This new understanding will guide new experimental and clinical trial designs.

## MATERIALS AND METHODS

### Animals

All experiments were approved by the University of Florida Institutional Animal Care and Use Committee. Experiments were performed on young (3-4 m.o.) male Sprague-Dawley rats (208A Colony, Envigo; IN, USA). Rats were housed in pairs at 24°C with a 12/12 light/dark cycle (lights on at 07:00/lights off at 19:00) and access to food and water *ad libitum*. All rats underwent a 14-day entrainment period prior to use in experiments. Sample size estimation was based on previous studies ^67,84,85^ and our extensive experience with this neurophysiological preparation.

### Spinal Tissue Sample Collection

Rats were euthanized during midrest (12:00) and midactive (00:00) time points. During the midactive phase (00:00) tissues were collected under a dim red light to minimize light cues. Rats were euthanized under isoflurane anesthesia (4%) via intracardiac perfusion of 1X phosphate-buffered saline (PBS) with a peristaltic pump (Masterflex, Cole-Palmer). Cervical spinal tissue containing the phrenic motor nucleus (C_3_-C_5_) was collected and transferred to a microtome (−26°C) to separate ventral from dorsal cervical spinal cord. Tissue was flash frozen in liquid nitrogen and stored at −80°C until use (< 3 weeks). The mean sample wet weight for ventral C_3_–C_5_ spinal cord was: mean (*M*) =□47.71 mg, standard error mean (SEM) = 1.44 mg.

### Spinal Adenosine and Serotonin Measurements

Tissue was homogenized in 1X PBS and 200 nM adenosine kinase inhibitor ABT 702 dihydrochloride (Tocris Bioscience) at a concentration of 100 mg tissue/ 1 mL PBS. Homogenized samples were then centrifuged at 10,000 x g for 10 minutes at 4°C. The supernatant is then assayed directly.

#### ADENOSINE ASSAY

The adenosine assay was performed using a coupled enzyme reaction according to the method described by the manufacturer (Cell BioLabs, Inc., REF # MET-5090) and adapted from Jagernaath, et al ^41^. Adenosine in ventral C_3_-C_5_ or standard is converted to xanthine and hydrogen peroxide through a series of enzymatic reactions. Horseradish peroxidase catalyzes the reaction between the resulting hydrogen peroxide and the Oxired probe to bind in a 1:1 ratio. The fluorescence was measured using a FlexStation 3 Multimode Plate Reader (Molecular Devices, San Jose, CA) with excitation at 560 nm and emission at 590 nm. All samples were assayed with and without adenosine deaminase to account for background. The data was analyzed by subtracting fluorescence values from background and normalizing to protein content. Intra-assay variability was 5.603%.

#### SEROTONIN ELISA

Spinal C_3_-C_5_ serotonin levels were measured using a rat serotonin ELISA (MyBioSource, REF # MBS166089) according to methods described by the manufacturer. Serotonin present in samples binds to rat serotonin antibody that has been pre-coated on plate wells. Biotinylated rat serotonin antibody binds to serotonin in the sample, and then bound by Streptavidin-HRP. After incubation, substrate solution is added, and color develops proportionately to the amount of rat serotonin. The reaction is terminated by addition of acidic stop solution and absorbance is measured at 450 nm. Intra-assay variability was 4.662%.

### Surgical Preparation

Terminal experiments were performed as previously described ^16,23,72,84^. Rats were induced with 3.0% isoflurane in oxygen in a plexiglass chamber, and transferred to a heated surgical table where anesthesia was continued (3.0% isoflurane; 60% oxygen, balance nitrogen). Body temperature was monitored with a rectal thermometer (Fisher Scientific, Pittsburgh, PA) and maintained between 37.0 ± 1.0°C throughout the experiment. A polyethylene catheter (I.D. 1.67 mm; PE 240; Intramedic, MD) was inserted into the trachea through a mid-line neck incision, and rats were mechanically ventilated (0.07mL/10g bw/ breath; 72 breaths/min; VentElite small animal ventilator; Harvard Apparatus, Holliston, MA, USA). End-tidal P_CO2_ (PET_CO2_) was monitored throughout the preparation and experimental protocol (Capnogard, Novametrix, Wallingford CT). The inspired CO_2_ fraction was adjusted as needed to keep end-tidal CO_2_ within physiological limits.

Anesthesia was then slowly converted intravenously to urethane through tail vein catheter (2.1g/kg at 6mL/h; 24 gauge, Surflo, Elkton, MD); while progressively decreasing the inspired isoflurane concentration 0.5% every 6 min. Anesthetic depth was assessed throughout conversion by toe-pinch withdrawal reflex; supplemental anesthetic was infused as required. Rats were then bilaterally vagotomized at the mid-cervical level to prevent phrenic nerve entrainment with the ventilator. The right femoral artery was isolated and cannulated with polyethylene tubing (I.D. 0.58 mm; PE 50; Intradermic, MD) to monitor blood pressure (TA-100 Transducer Amplifier, CWE, Inc.) and sample blood gases (PaO_2_, PaCO_2_, acid-base balance, pH and hemoglobin using a blood gas analyzer (ABL 90 Flex, Radiometer, Copenhagen, Denmark). Once urethane infusion was complete, fluids were administered intravenously to maintain acid-base balance (1.5 mL/h; 1:4 of 8.4% Na_2_CO_3_ in standard lactated Ringer’s solution). Rats were also paralyzed with pancronium bromide (2 mg/kg; Sigma-Aldrich, St. Louis, MO).

### SERIES I: Micro-Optode Measurements for Spinal PtO_2_

A dorsal midline incision was made from the base of the skull to the fifth cervical segment. Muscle layers were retracted to expose the C_3_-C_5_ vertebrae. A laminectomy was performed at C_4_ vertebrae. A longitudinal cut was made in the dura mater to facilitate insertion of a 50 μm oxygen micro-optode mounted on a micromanipulator (Unisense; Aarhus, Denmark). The tip of the optical sensor was coated with a fluorophore that, when excited with 610 nm red light pulses, emits 780 nm infrared light that varies with intensity inversely proportional to the oxygen partial pressure. The micro-optode was placed on the left side 1 mm lateral to midline at ~1.5 mm depth to measure PtO_2_ near/in the phrenic motor nucleus, as previously described ^48^. Coordinates were obtained from previous studies ^86^.

Raw signals were acquired at 1 Hz and converted to mmHg using a two-point calibration. Saline was warmed to 37 °C and aerated with 21% oxygen over 40 min for the first calibration point (i.e. normoxia). Sodium hydroxide (0.1 M) and sodium ascorbate (0.1 M) were then added in warmed saline to create an anoxic solution for the second calibration point (i.e. zero oxygen). Measurements were taken about approximately 45 minutes after probe placement. Rats were exposed to 5-min of 12-14% O_2_ or 1-min of 8-10% O_2_ (n = 4 each group; PaO_2_ = 40-50 mmHg). Calibration was verified *in vivo* by PtO_2_ values reaching 0–1 mmHg several minutes after death from a urethane overdose (not shown).

### SERIES II: Adenosine and Inosine Probe Measurements for Changes in Spinal Extracellular Adenosine

In a separate set of rats (n = 3) changes in extracellular adenosine concentrations (ΔADO) during 5-min of 13% O_2_ and 1-min of 9% O_2_ (PaO_2_ = 40-50 mmHg). Following the same surgical procedures and probe placement coordinates described in ‘**SERIES I’** experiments, extracellular ΔADO concentrations were measured by differential enzymatic detection method using adenosine and inosine microbiosensors (Zimmer-Peacock, UK) positioned 1 mm lateral to midline between C_3_ and C_4_ at ~1.5 mm depth. Microbiosensors were approximately 3-4 mm apart from each other on the same side of the cervical spinal cord. The signals were acquired at 1 Hz and converted to concentrations determined using a three-point calibration. Measurements were taken about 30-45 minutes after probe insertion. The functionality of probes was confirmed at the end of each experiment.

### SERIES III: Neurophysiological Experiments

A laminectomy was performed at the C_2_ vertebrae for intrathecal drug delivery. A small hole was then cut in the dura near the junction of C_2_ and C_3_ spinal segment, and a flexible silicone catheter (O.D. 0.6 mm; Access Technologies) was fed through to the caudal end of C_3_. A 50 μL Hamilton syringe containing drug (*see* ‘Drugs and Vehicles’ Section) was attached to the catheter for drug delivery. To prevent off target effects of pharmacological manipulation, we used intrathecal injection of drugs directly at C_4_ instead of systemic administration to limit unintended drug distribution. For example, in an anatomically separated structure, respiratory LTF in hypoglossal (XII) motor output (brainstem structure rostral the phrenic nerve) has been measured simultaneously in previous reports from our lab and found hypoglossal motor output to be unaffected by intrathecal drug delivery directly at C_4_ ^14,23^.

### Drugs and Vehicles

Drugs used throughout studies include MSX-3 (A_2A_ receptor antagonist; #M3568; Millipore Sigma), ketanserin tartrate (5-HT_2A/C_ receptor antagonist; #090850; FisherSci) and istradefylline (A_2A_ receptor antagonist; #51470R; FisherSci). Upon arrival, all drugs were initially dissolved in 100% DMSO or 0.9% saline based on solubility and manufacturer information. Aliquots of the stock solution were kept frozen in −20°C. On the day of the experiments, all drugs were diluted in sterile 0.9% saline to achieve the desired final concentration. The final DMSO-saline ratios were determined by the solubility of the compound; a final concentration of 10% DMSO was sufficient at maintaining all drugs dissolved in the vehicle solution.

Most drugs were dissolved to final effective concentrations as previously determined in our laboratory via dose-response studies ^15^. Dose-response studies determined *1*) effective doses for each agonist in inducing phrenic motor facilitation, *2*) efficacy of each agonist in the presence of its selective antagonist, and *3*) relative selectivity for each agonist in the presence of other receptor subtype antagonists. Based on these (and other) previous findings, intrathecal drug doses were as follows: 10 μM MSX-3 (130 ng/kg, 12 μL) ^16^, 1 mM istradefylline (10 μg/kg, 12-15 μL), and 500 μM ketanserin tartrate (10 μg/kg, 18 μL).

### Electrophysiological Recordings

Using a dorsal approach, the left phrenic nerve was isolated, cut distally, and de-sheathed. Custom suction recording electrodes filled with 0.9% saline were then placed in the saline-filled phrenic pocket and the nerve was suctioned up with a 60 mL syringe to record respiratory neural activity.

Nerve activity was amplified (10K, A-M systems, Everett, WA), filtered (bandpass 300-5,000 Hz), integrated (time constant, 50 ms), digitized (CED 1401, Cambridge Electronic Design, UK), and analyzed using Spike2 software (CED, version 8.20). Inspiratory phrenic activity served as an index of respiratory motor output.

### Experimental Protocols

At least 45 min after conversion to urethane anesthesia, apneic and recruitment CO_2_ thresholds of respiratory nerve activity were determined by: 1) lowering inspired CO_2_ levels or 2) increasing ventilation rate until rhythmic respiratory nerve activity ceased. After 1 min, inspired CO_2_ levels was slowly increased until respiratory rhythmic respiratory nerve bursts resumed. Baseline conditions were established at P_ETCO_2__ 2 mmHg above the recruitment threshold. Blood samples were taken during baseline to document baseline blood gas levels during stable nerve activity. Arterial PCO_2_ was maintained isocapnic (± 1.5 mmHg) with respect to baseline blood gas values by actively manipulating inspired CO_2_ concentration and/or ventilation rate. Baseline oxygen levels (60% oxygen, balance nitrogen and carbon dioxide; PaO_2_ > 150 mmHg) were maintained for the duration of the experiments, except for hypoxic challenges (PaO_2_ = 40-55 mmHg). At the end of the protocol, a maximum chemoreceptor challenge (10% O_2_, 83% N_2_, and 7% CO_2_) was administered for 3 minutes to ensure phrenic nerve amplitude was not reaching a ceiling effect during the mAIH protocol (data in *Supplement*). Rats were sacrificed by urethane overdose.

### Statistical Analyses

Measurements of peak integrated amplitude and inspiratory phrenic burst frequency (bursts/min) were assessed in 1 min bins immediately before each blood sample at: baseline, last minute of first hypoxic episode (short-term hypoxic response; STHR) 30, 60, and 90 min post-AIH, and during the final minute of the maximum chemoreceptor challenge (**Table S3**). Measurements were also made at equivalent time points in time-matched control experiments. Nerve burst amplitudes were normalized by subtracting from the baseline value, then dividing by the baseline and reported as percentage change from baseline. Burst frequencies were also normalized to baseline, but expressed as an absolute difference in bursts per minute. All statistical comparisons between treatment groups for nerve amplitude (baseline and 90 min time points) were made using a two or three-way ANOVA with a repeated measures design. All individual comparisons were made using Tukey *post-hoc* test.

Comparisons for mean arterial pressures, arterial P_CO_2__ and P_O_2__ (**Tables S1** and **S2**) and respiratory frequency (**Table S4**) were made at baseline, hypoxia episode 1 (and hypoxia episodes 8 and 15 for the 15×1 mAIH protocol), 30, 60 and 90 min post-AIH using two-way ANOVA to determine if there was an effect of time of day on baseline values; two-way mixed effects ANOVA was then used to test if there was an effect of protocol with drug pretreatment throughout the protocol. Comparisons in spinal adenosine and serotonin levels during midrest and midactive time points were made with an unpaired *t*-test. The spinal PtO_2_ signal was smoothed and analyzed as one-minute averages before AIH, during the nadir of the first hypoxic episode, and at the peak during reoxygenation following the first hypoxic episode. Unpaired *t*-tests were used to compare measurements from PtO_2_ and ΔADO experiments during 5 min and 1 min (PaO_2_ = 40-50 mmHg) hypoxic exposures.

Three-way repeated measure ANOVA and mixed effect ANOVA designs were calculated using JMP Pro (version 16.1.0; SAS Institute, Inc. Cary, NC, USA). All other statistics were analyzed in SigmaPlot (version 14.0.0.124; Systat Software Inc., San Jose, CA, USA). Differences between groups were considered significant if p < 0.050. Data are presented as mean ± standard error mean (SEM).

## Acknowledgements

The authors would like to thank J. Oberto and C. Lurk for their technical assistance.

## Funding

This work was supported by NIH R01HL148030 (GSM), NIH R01HL149800 (GSM), and NIH T32HL134621-5 (ABM, MNK), and the UF Brain Spinal Cord Research Trust Fund.

## Author contributions

Conceptualization: *ABM, MNK, GSM*. Data curation: *ABM, YBS, RRP, GSM*. Formal analysis: *ABM, GSM*. Funding acquisition: *GSM*. Investigation: *ABM, YBS, MNK, RRP, GSM*. Methodology: *ABM, GSM*. Supervision: *GSM*. Writing (original draft): *ABM, GSM*. Manuscript revisions: *ABM, YBS, MNK, RRP, GSM*.

## Competing interests

The authors declare no competing interests.

## Data and Materials Availability

All data needed to evaluate the conclusions presented are available in the paper and/or the Supplementary Materials.

## Supplementary Materials

Supplementary text

Figures S1-S4

Tables S1-S4

## Notes

### Competing Interest Statement

The authors have declared no competing interest.

## REFERENCES

1. Mitchell, G.S. & Johnson, S.M. Neuroplasticity in respiratory motor control. J Appl Physiol (1985) 94, 358–374 (2003).

2. Mitchell, G.S. & Baker, T.L. Respiratory neuroplasticity: Mechanisms and translational implications of phrenic motor plasticity. Handb Clin Neurol 188, 409–432 (2022).

3. Vose, A.K., et al. Therapeutic acute intermittent hypoxia: A translational roadmap for spinal cord injury and neuromuscular disease. Exp Neurol 347, 113891 (2022).

4. Gonzalez-Rothi, E.J., et al. Intermittent hypoxia and neurorehabilitation. J Appl Physiol (1985) 119, 1455–1465 (2015).

5. Dale, E.A., Ben Mabrouk, F. & Mitchell, G.S. Unexpected benefits of intermittent hypoxia: enhanced respiratory and nonrespiratory motor function. Physiology (Bethesda) 29, 39–48 (2014).

6. Sajjadi, E., et al. Acute intermittent hypoxia and respiratory muscle recruitment in people with amyotrophic lateral sclerosis: A preliminary study. Exp Neurol 347, 113890 (2022).

7. Millhorn, D.E., Eldridge, F.L. & Waldrop, T.G. Prolonged stimulation of respiration by a new central neural mechanism. Respir Physiol 41, 87–103 (1980).

8. Hayashi, F., Coles, S.K., Bach, K.B., Mitchell, G.S. & McCrimmon, D.R. Time-dependent phrenic nerve responses to carotid afferent activation: intact vs. decerebellate rats. Am J Physiol 265, R811–819 (1993).

9. Fregosi, R.F. & Mitchell, G.S. Long-term facilitation of inspiratory intercostal nerve activity following carotid sinus nerve stimulation in cats. J Physiol 477 (Pt 3), 469–479 (1994).

10. Feldman, J.L., Mitchell, G.S. & Nattie, E.E. Breathing: rhythmicity, plasticity, chemosensitivity. Annu Rev Neurosci 26, 239–266 (2003).

11. Mitchell, G.S., et al. Invited review: Intermittent hypoxia and respiratory plasticity. J Appl Physiol (1985) 90, 2466–2475 (2001).

12. Millhorn, D.E., Eldridge, F.L. & Waldrop, T.G. Prolonged stimulation of respiration by endogenous central serotonin. Respir Physiol 42, 171–188 (1980).

13. Bach, K.B. & Mitchell, G.S. Hypoxia-induced long-term facilitation of respiratory activity is serotonin dependent. Respir Physiol 104, 251–260 (1996).

14. Baker-Herman, T.L. & Mitchell, G.S. Phrenic long-term facilitation requires spinal serotonin receptor activation and protein synthesis. J Neurosci 22, 6239–6246 (2002).

15. Tadjalli, A. & Mitchell, G.S. Cervical Spinal 5-HT2A and 5-HT2B receptors are both necessary for moderate acute intermittent hypoxia-induced phrenic long-term facilitation. J Appl Physiol (1985) (2019).

16. Nichols, N.L., Dale, E.A. & Mitchell, G.S. Severe acute intermittent hypoxia elicits phrenic longterm facilitation by a novel adenosine-dependent mechanism. J Appl Physiol (1985) 112, 1678–1688 (2012).

17. Seven, Y.B., et al. Phrenic motor neuron adenosine 2A receptors elicit phrenic motor facilitation. J Physiol 596, 1501–1512 (2018).

18. Dale-Nagle EA, H.M., MacFarlane PM, Mitchell GS. Multiple pathways to long-lasting phrenic motor facilitation. Adv Exp Med Biol 669, 225–230 (2010).

19. Devinney, M.J., Huxtable, A.G., Nichols, N.L. & Mitchell, G.S. Hypoxia-induced phrenic longterm facilitation: emergent properties. Ann N Y Acad Sci 1279, 143–153 (2013).

20. Fields, D.P. & Mitchell, G.S. Divergent cAMP signaling differentially regulates serotonin-induced spinal motor plasticity. Neuropharmacology 113, 82–88 (2017).

21. Perim, R.R. & Mitchell, G.S. Circulatory control of phrenic motor plasticity. Respir Physiol Neurobiol 265, 19–23 (2019).

22. Devinney, M.J., Nichols, N.L. & Mitchell, G.S. Sustained Hypoxia Elicits Competing Spinal Mechanisms of Phrenic Motor Facilitation. J Neurosci 36, 7877–7885 (2016).

23. Hoffman, M.S., Golder, F.J., Mahamed, S. & Mitchell, G.S. Spinal adenosine A2(A) receptor inhibition enhances phrenic long term facilitation following acute intermittent hypoxia. J Physiol 588, 255–266 (2010).

24. Porkka-Heiskanen, T., Strecker, R.E. & McCarley, R.W. Brain site-specificity of extracellular adenosine concentration changes during sleep deprivation and spontaneous sleep: an in vivo microdialysis study. Neuroscience 99, 507–517 (2000).

25. Bjorness, T.E. & Greene, R.W. Adenosine and sleep. Curr Neuropharmacol 7, 238–245 (2009).

26. Huang, Z.L., Urade, Y. & Hayaishi, O. The role of adenosine in the regulation of sleep. Curr Top Med Chem 11, 1047–1057 (2011).

27. Tester, N.J., et al. Long-term facilitation of ventilation in humans with chronic spinal cord injury. Am J Respir Crit Care Med 189, 57–65 (2014).

28. Tester, N.J., Fuller, D.D. & Mateika, J.H. Ventilatory long-term facilitation in humans. Am J Respir Crit Care Med 189, 1009–1010 (2014).

29. Winn, H.R., Rubio, R. & Berne, R.M. Brain adenosine concentration during hypoxia in rats. Am J Physiol 241, H235–242 (1981).

30. Martin, E.D., et al. Adenosine released by astrocytes contributes to hypoxia-induced modulation of synaptic transmission. Glia 55, 36–45 (2007).

31. Bannon, N.M., Chistiakova, M., Chen, J.Y., Bazhenov, M. & Volgushev, M. Adenosine Shifts Plasticity Regimes between Associative and Homeostatic by Modulating Heterosynaptic Changes. J Neurosci 37, 1439–1452 (2017).

32. Porkka-Heiskanen, T., et al. Adenosine: a mediator of the sleep-inducing effects of prolonged wakefulness. Science 276, 1265–1268 (1997).

33. Chaudhury, D., Wang, L.M. & Colwell, C.S. Circadian regulation of hippocampal long-term potentiation. J Biol Rhythms 20, 225–236 (2005).

34. Seibt, J. & Frank, M.G. Primed to Sleep: The Dynamics of Synaptic Plasticity Across Brain States. Front Syst Neurosci 13, 2 (2019).

35. Gerstner, J.R. & Yin, J.C. Circadian rhythms and memory formation. Nat Rev Neurosci 11, 577-588 (2010).

36. Bellesi, M. & de Vivo, L. Structural synaptic plasticity across sleep and wake. Curr Opin Physiol 15, 74–81 (2020).

37. Huston, J.P., et al. Extracellular adenosine levels in neostriatum and hippocampus during rest and activity periods of rats. Neuroscience 73, 99–107 (1996).

38. Liu, D.K., Horner, R.L. & Wojtowicz, J.M. Time of day determines modulation of synaptic transmission by adenosine in the rat hippocampal slices. Neurosci Lett 282, 200–202 (2000).

39. Rex, C.S., et al. Different Rho GTPase-dependent signaling pathways initiate sequential steps in the consolidation of long-term potentiation. J Cell Biol 186, 85–97 (2009).

40. Kelly, M.N., et al. Circadian clock genes and respiratory neuroplasticity genes oscillate in the phrenic motor system. Am J Physiol Regul Integr Comp Physiol 318, R1058–R1067 (2020).

41. Jagannath, A., et al. Adenosine integrates light and sleep signalling for the regulation of circadian timing in mice. Nat Commun 12, 2113 (2021).

42. Zhang, Z., Wang, H.J., Wang, D.R., Qu, W.M. & Huang, Z.L. Red light at intensities above 10 lx alters sleep-wake behavior in mice. Light Sci Appl 6, e16231 (2017).

43. Jimenez-Zarate, B.S., et al. Day-Night Variations in the Concentration of Neurotransmitters in the Rat Lumbar Spinal Cord. J Circadian Rhythms 19, 9 (2021).

44. Perim, R.R., El-Chami, M., Gonzalez-Rothi, E.J. & Mitchell, G.S. Baseline Arterial CO2 Pressure Regulates Acute Intermittent Hypoxia-Induced Phrenic Long-Term Facilitation in Rats. Front Physiol 12, 573385 (2021).

45. Harada, Y., Kuno, M. & Wang, Y.Z. Differential effects of carbon dioxide and pH on central chemoreceptors in the rat in vitro. J Physiol 368, 679–693 (1985).

46. Mateika, J.H. & Fregosi, R.F. Long-term facilitation of upper airway muscle activities in vagotomized and vagally intact cats. J Appl Physiol (1985) 82, 419–425 (1997).

47. Kumar, P. & Prabhakar, N.R. Peripheral chemoreceptors: function and plasticity of the carotid body. Compr Physiol 2, 141–219 (2012).

48. Perim, R.R., Gonzalez-Rothi, E.J. & Mitchell, G.S. Cervical spinal injury compromises caudal spinal tissue oxygenation and undermines acute intermittent hypoxia-induced phrenic long-term facilitation. Exp Neurol 342, 113726 (2021).

49. Hoffman, M.S., Nichols, N.L., Macfarlane, P.M. & Mitchell, G.S. Phrenic long-term facilitation after acute intermittent hypoxia requires spinal ERK activation but not TrkB synthesis. J Appl Physiol (1985) 113, 1184–1193 (2012).

50. Dale, E.A., Fields, D.P., Devinney, M.J. & Mitchell, G.S. Phrenic motor neuron TrkB expression is necessary for acute intermittent hypoxia-induced phrenic long-term facilitation. Exp Neurol 287, 130–136 (2017).

51. Baker-Herman, T.L., et al. BDNF is necessary and sufficient for spinal respiratory plasticity following intermittent hypoxia. Nat Neurosci 7, 48–55 (2004).

52. Wilkerson, J.E. & Mitchell, G.S. Daily intermittent hypoxia augments spinal BDNF levels, ERK phosphorylation and respiratory long-term facilitation. Exp Neurol 217, 116–123 (2009).

53. Marmigere, F., Givalois, L., Rage, F., Arancibia, S. & Tapia-Arancibia, L. Rapid induction of BDNF expression in the hippocampus during immobilization stress challenge in adult rats. Hippocampus 13, 646–655 (2003).

54. Golder, F.J., et al. Spinal adenosine A2a receptor activation elicits long-lasting phrenic motor facilitation. J Neurosci 28, 2033–2042 (2008).

55. Powell, F.L., Milsom, W.K. & Mitchell, G.S. Time domains of the hypoxic ventilatory response. Respir Physiol 112, 123–134 (1998).

56. Pamenter, M.E. & Powell, F.L. Time Domains of the Hypoxic Ventilatory Response and Their Molecular Basis. Compr Physiol 6, 1345–1385 (2016).

57. Hayes, H.B., et al. Daily intermittent hypoxia enhances walking after chronic spinal cord injury: a randomized trial. Neurology 82, 104–113 (2014).

58. Trumbower, R.D., Hayes, H.B., Mitchell, G.S., Wolf, S.L. & Stahl, V.A. Effects of acute intermittent hypoxia on hand use after spinal cord trauma: A preliminary study. Neurology 89, 1904–1907 (2017).

59. Trumbower, R.D., et al. Caffeine enhances intermittent hypoxia-induced gains in walking function for people with chronic spinal cord injury. J Neurotrauma (2022).

60. Navarrete-Opazo, A., Dougherty, B.J. & Mitchell, G.S. Enhanced recovery of breathing capacity from combined adenosine 2A receptor inhibition and daily acute intermittent hypoxia after chronic cervical spinal injury. Exp Neurol 287, 93–101 (2017).

61. Kell, C.A. & Stehle, J.H. Just the two of us: melatonin and adenosine in rodent pituitary function. Ann Med 37, 105–120 (2005).

62. Imura, Y., et al. Microglia release ATP by exocytosis. Glia 61, 1320–1330 (2013).

63. Sebastiao, A.M., Cunha, R.A., de Mendonca, A. & Ribeiro, J.A. Modification of adenosine modulation of synaptic transmission in the hippocampus of aged rats. Br J Pharmacol 131, 1629–1634 (2000).

64. Costenla, A.R., et al. Enhanced role of adenosine A(2A) receptors in the modulation of LTP in the rat hippocampus upon ageing. Eur J Neurosci 34, 12–21 (2011).

65. Canas, P.M., Duarte, J.M., Rodrigues, R.J., Kofalvi, A. & Cunha, R.A. Modification upon aging of the density of presynaptic modulation systems in the hippocampus. Neurobiol Aging 30, 1877–1884 (2009).

66. Huxtable, A.G., Smith, S.M., Vinit, S., Watters, J.J. & Mitchell, G.S. Systemic LPS induces spinal inflammatory gene expression and impairs phrenic long-term facilitation following acute intermittent hypoxia. J Appl Physiol (1985) 114, 879–887 (2013).

67. Behan, M., Zabka, A.G. & Mitchell, G.S. Age and gender effects on serotonin-dependent plasticity in respiratory motor control. Respir Physiol Neurobiol 131, 65–77 (2002).

68. Martinez, A., et al. TrkB and TrkC signaling are required for maturation and synaptogenesis of hippocampal connections. J Neurosci 18, 7336–7350 (1998).

69. Miao, L., et al. mTORC1 is necessary but mTORC2 and GSK3beta are inhibitory for AKT3-induced axon regeneration in the central nervous system. Elife 5, e14908 (2016).

70. Perim, R.R., Fields, D.P. & Mitchell, G.S. Spinal AMP kinase activity differentially regulates phrenic motor plasticity. J Appl Physiol (1985) 128, 523–533 (2020).

71. Nichols, N.L. & Mitchell, G.S. Mechanisms of severe acute intermittent hypoxia-induced phrenic long-term facilitation. J Neurophysiol 125, 1146–1156 (2021).

72. Tadjalli, A., Seven, Y.B., Perim, R.R. & Mitchell, G.S. Systemic inflammation suppresses spinal respiratory motor plasticity via mechanisms that require serine/threonine protein phosphatase activity. J Neuroinflammation 18, 28 (2021).

73. Dale, E.A., Satriotomo, I. & Mitchell, G.S. Cervical spinal erythropoietin induces phrenic motor facilitation via extracellular signal-regulated protein kinase and Akt signaling. J Neurosci 32, 5973–5983 (2012).

74. Mohawk, J.A., Pargament, J.M. & Lee, T.M. Circadian dependence of corticosterone release to light exposure in the rat. Physiol Behav 92, 800–806 (2007).

75. Dauchy, R.T., et al. The influence of red light exposure at night on circadian metabolism and physiology in Sprague-Dawley rats. J Am Assoc Lab Anim Sci 54, 40–50 (2015).

76. Terada, J., et al. Ventilatory long-term facilitation in mice can be observed during both sleep and wake periods and depends on orexin. J Appl Physiol (1985) 104, 499–507 (2008).

77. Zabka, A.G., Behan, M. & Mitchell, G.S. Selected contribution: Time-dependent hypoxic respiratory responses in female rats are influenced by age and by the estrus cycle. J Appl Physiol (1985) 91, 2831–2838 (2001).

78. Behan, M., Zabka, A.G., Thomas, C.F. & Mitchell, G.S. Sex steroid hormones and the neural control of breathing. Respir Physiol Neurobiol 136, 249–263 (2003).

79. Poulsen, R.C., Warman, G.R., Sleigh, J., Ludin, N.M. & Cheeseman, J.F. How does general anaesthesia affect the circadian clock? Sleep Med Rev 37, 35–44 (2018).

80. Luo, M., Song, B. & Zhu, J. Sleep Disturbances After General Anesthesia: Current Perspectives. Front Neurol 11, 629 (2020).

81. Silver, N.R.G., Ward-Flanagan, R. & Dickson, C.T. Long-term stability of physiological signals within fluctuations of brain state under urethane anesthesia. PLoS One 16, e0258939 (2021).

82. Terada, J. & Mitchell, G.S. Diaphragm long-term facilitation following acute intermittent hypoxia during wakefulness and sleep. J Appl Physiol (1985) 110, 1299–1310 (2011).

83. Nakamura, A., et al. Sleep state dependence of ventilatory long-term facilitation following acute intermittent hypoxia in Lewis rats. J Appl Physiol (1985) 109, 323–331 (2010).

84. Zabka, A.G., Behan, M. & Mitchell, G.S. Long term facilitation of respiratory motor output decreases with age in male rats. J Physiol 531, 509–514 (2001).

85. Zabka, A.G., Mitchell, G.S. & Behan, M. Ageing and gonadectomy have similar effects on hypoglossal long-term facilitation in male Fischer rats. J Physiol 563, 557–568 (2005).

86. McGuire, M., Zhang, Y., White, D.P. & Ling, L. Phrenic long-term facilitation requires NMDA receptors in the phrenic motonucleus in rats. J Physiol 567, 599–611 (2005).

