## Supplemental Materials for "Daily fluctuations in spinal adenosine determine mechanisms of respiratory motor plasticity"

**This PDF file includes:**

Supplementary Text

Figs. S1 to S4

Tables S1 to S4

Supplementary Text

### *Spinal adenosine 2A receptor protein ELISA*

Ventral spinal C3-C5 segments were harvested and isolated as described in ***Materials and Methods*** section of the main text. Adenosine 2A receptor protein levels were measured using a rat ADORA2A/Adenosine A2A Receptor (Custom ELISA) ELISA kit (LS Biosciences, REF # LS-F34417) according to methods described by the manufacturer.

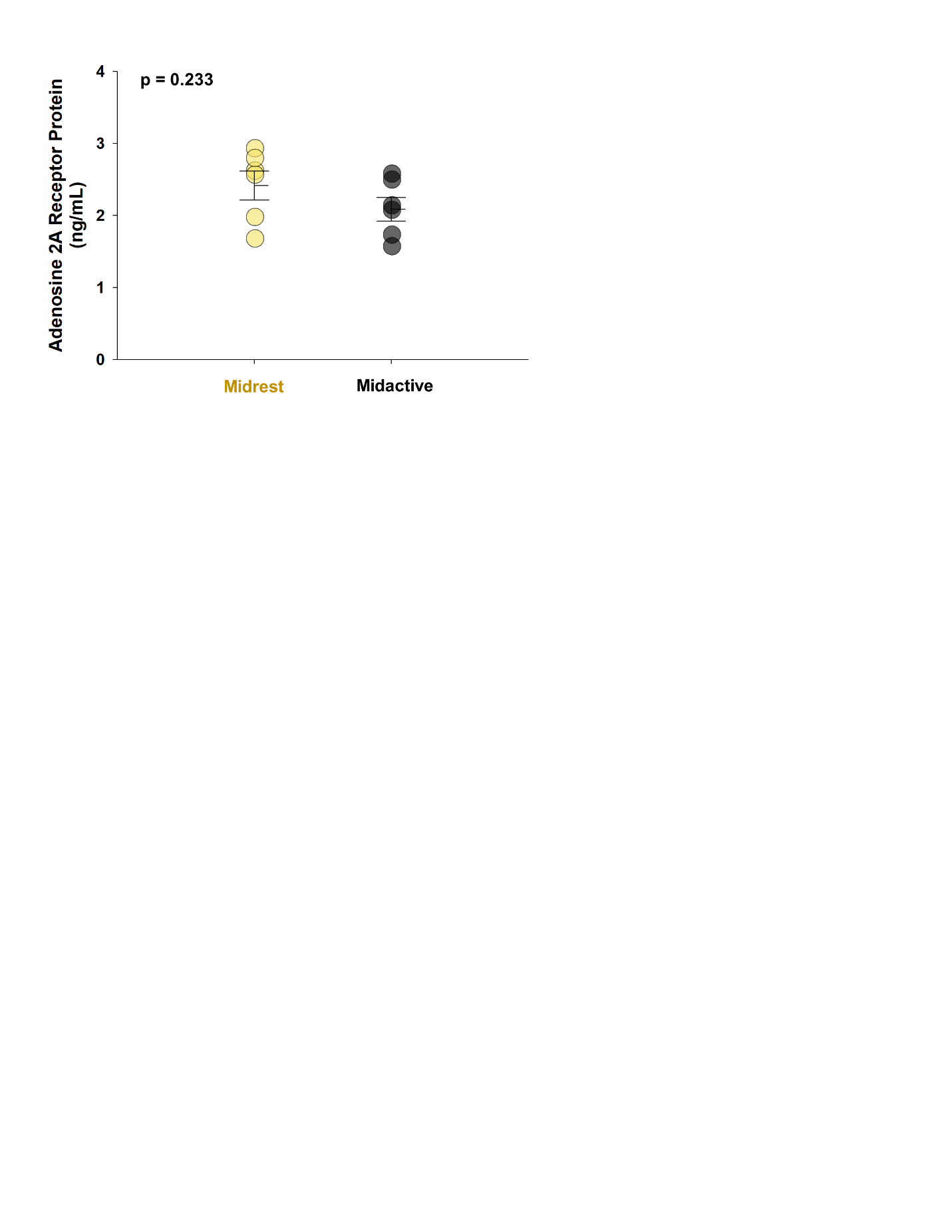

Figure S1. Adenosine 2A receptor protein in ventral cervical spinal homogenates are not different between midrest (noon) or midactive (midnight) time points. Adenosine 2A receptor protein concentrations in ventral C3-C5 tissue homogenates was similar in tissue harvested during midrest (12 PM) *versus* midactive (12 AM) phase (n = 6 each group; unpaired *t*-test, p = 0.233). Bars show mean ± SEM.

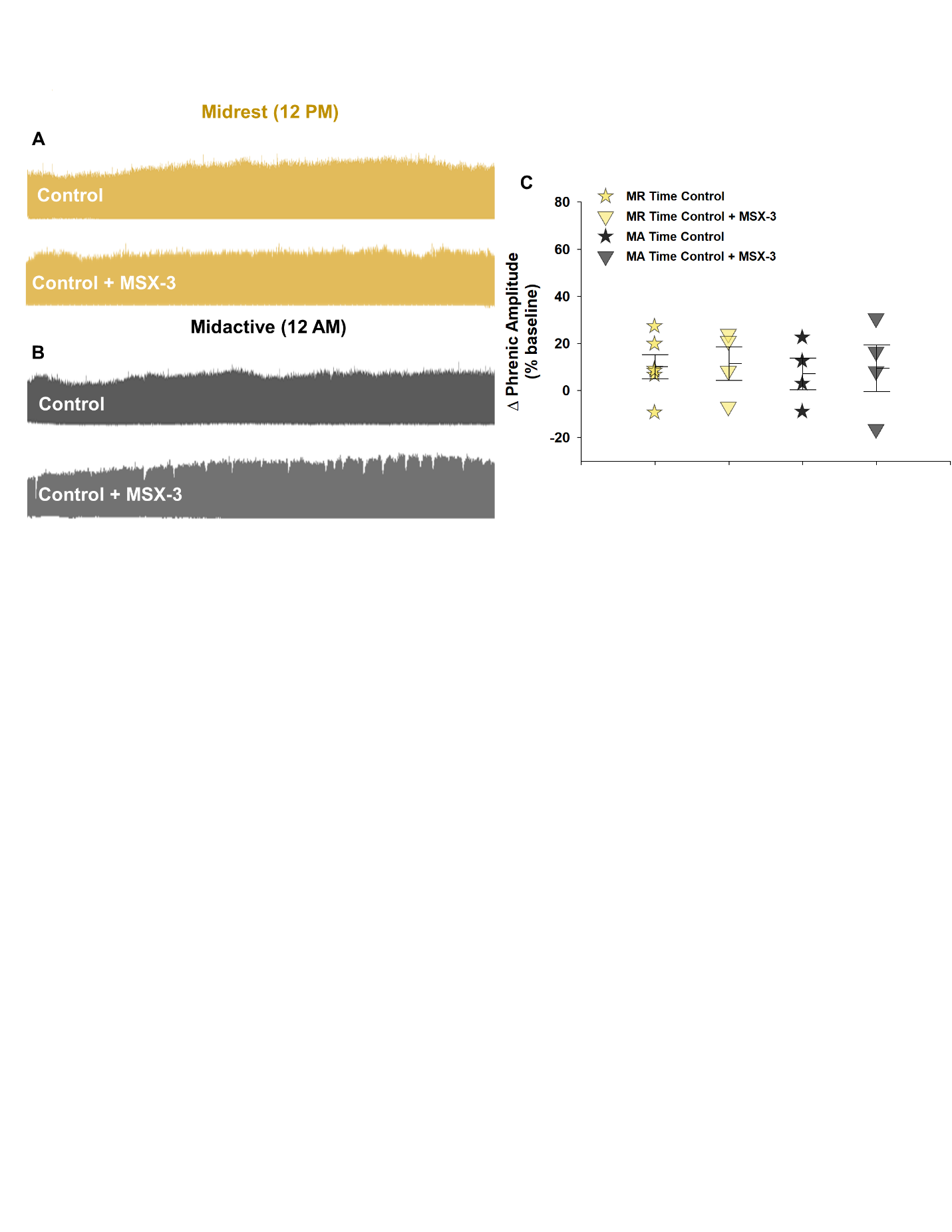

Figure S2. MSX-3 does not affect basal phrenic nerve activity. Representative neurograms during midrest (12 PM; A) or midactive (12 AM; B) time points without (*top*) or with intrathecally-administered MSX-3 (*bottom*). There is no effect of time of day or drug pretreatment on phrenic nerve activity 120 min after baseline (n = 4-6 each group; F_1,14_ = 0.215, p = 0.650; three-way RM ANOVA); C). Bars show mean ± SEM. MR, midrest; MA, midactive.

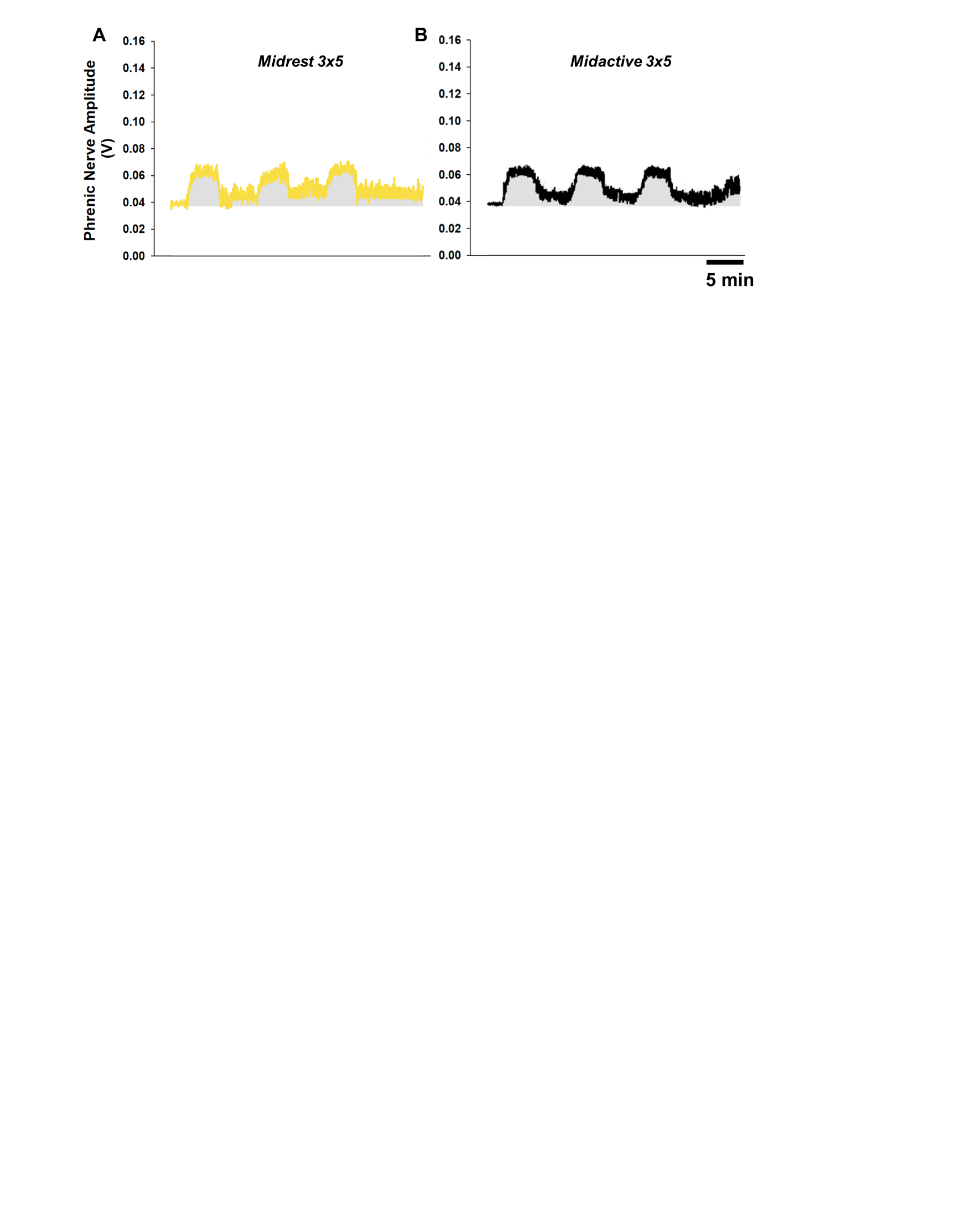

Figure S3. The 3x5 mAIH protocol does not exhibit pLTF “building” during midrest or midactive time points. Average traces of the 3x5 mAIH presented during midrest (n = 7; A) and midactive (n = 7; B) time points. There was no significant effect (one-way ANOVA; repeated measures) of the 3x5 mAIH protocol on phrenic nerve amplitude in the inter-hypoxic interval period (Midrest: F_1,6_ = 2.114; p = 0.196; Midactive: F_1,6_ = 1.341, p = 0.291).

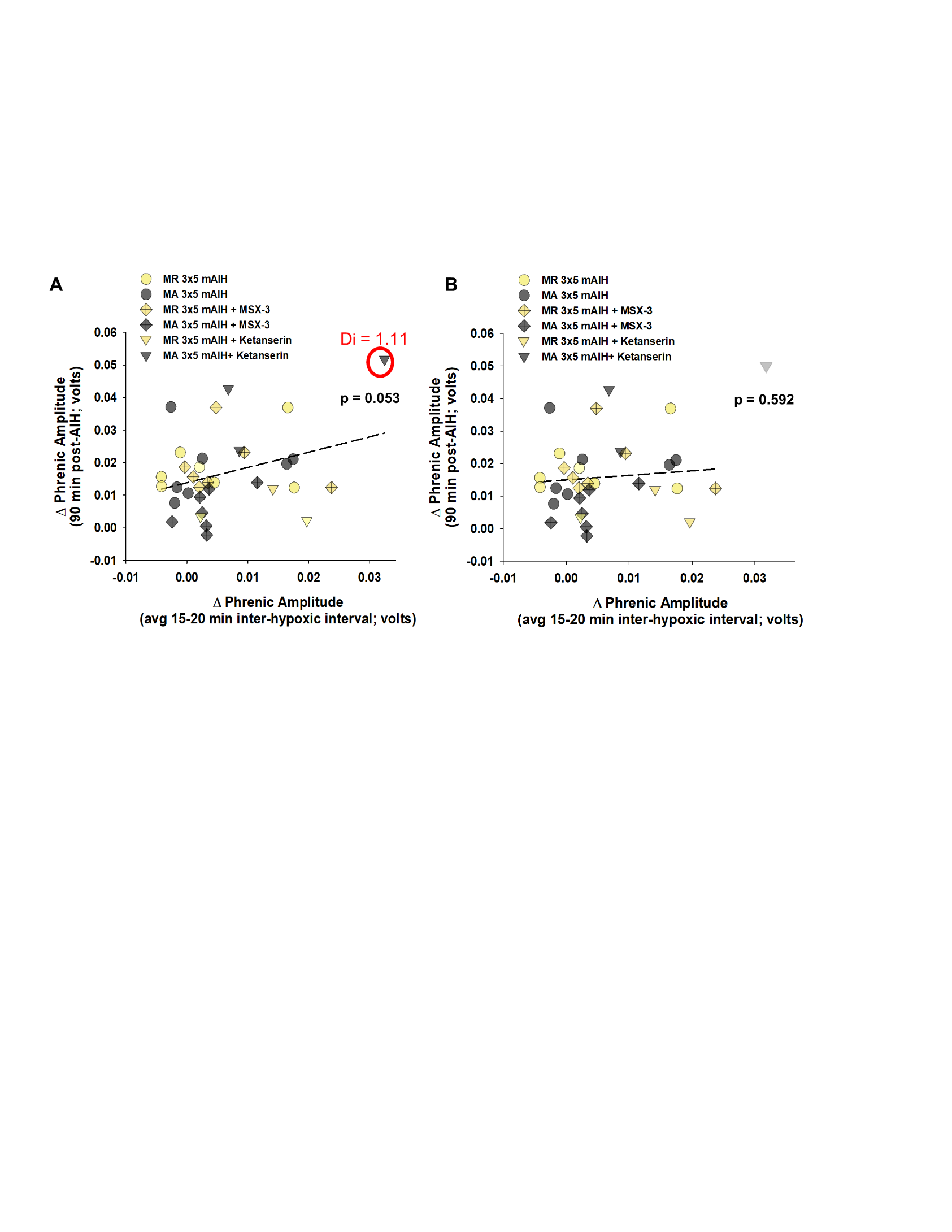

Figure S4. With longer hypoxic durations, phrenic nerve amplitude during inter-hypoxic intervals does not significantly correlate with pLTF 90 min post-AIH. Linear regression of all 3x5 mAIH midrest and midactive groups does not reveal a significant correlation between the change in phrenic nerve amplitude during the second inter-hypoxic interval and pLTF magnitude 90 min post-AIH. In A, the marginally significant correlation (r = 0.33; p = 0.053) is due to a single outlier. Cook’s Distance (Di) indicates an influential data point (*i.e.*, significant outlier) that is circled in red (A); when this outlier is removed (B) from the regression model, r value is reduced to 0.10, demonstrating the lack of any significant correlation (p = 0.59). MR, midrest; MA, midactive.

Table S1. Arterial PCO2, PO2, and MAP during baseline and 30, 60 and 90 min post-AIH

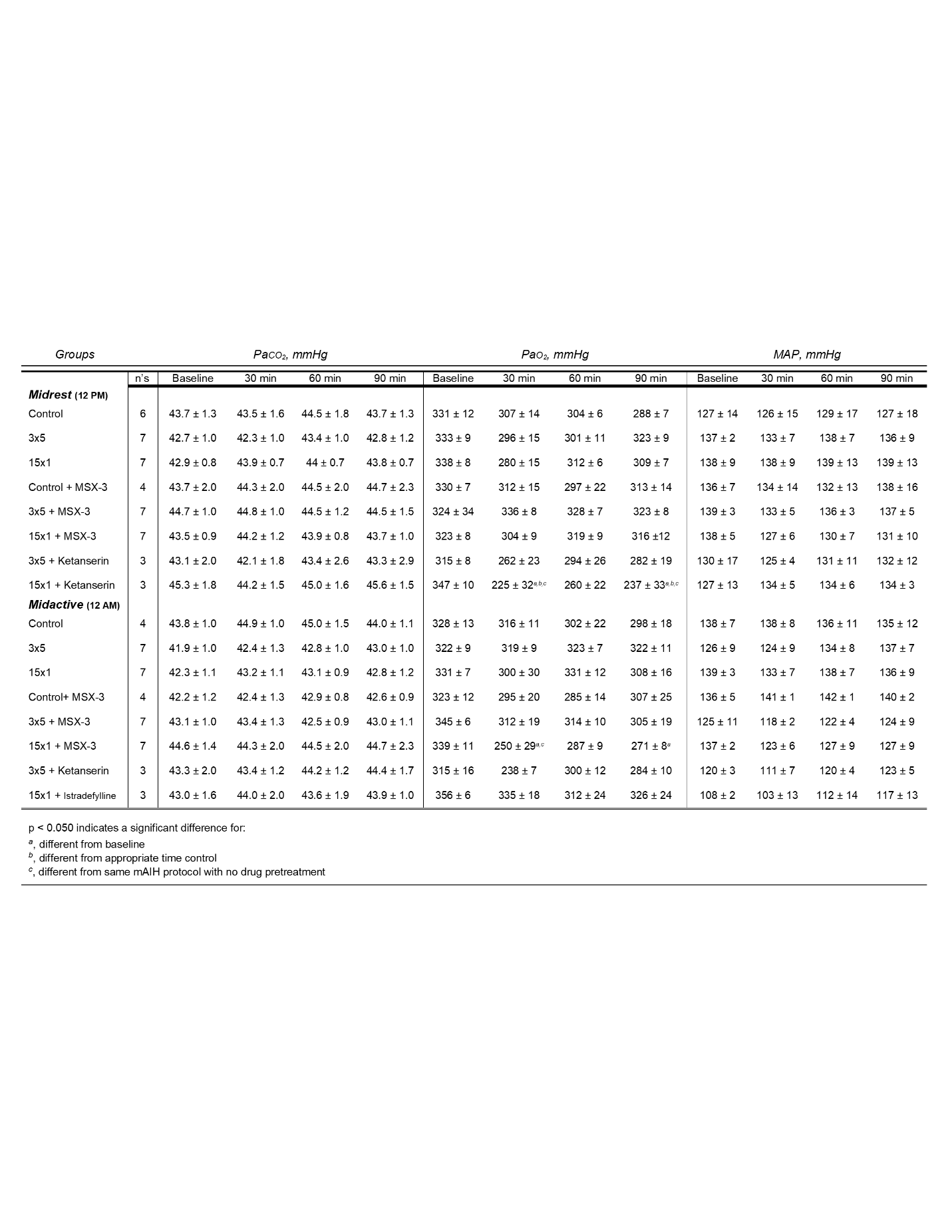

Two-way ANOVA was used to analyze blood gases and MAP. There was no effect of time of day or drug pre-treatment on baseline PaCO2 (F_6,79_ = 0.143, p = 0.990), PaO2 (F_6,79_ = 0.604, p = 0.727) or MAP (F_6,79_ = 1.508, p = 0.186). A two-way mixed effects ANOVA followed by Tukey *post-hoc* test was utilized to determine effect of protocol with drug pretreatment on PaCO2, PaO2, and MAP 30, 60, and 90 min post-mAIH during midrest or midactive phase. MAP, mean arterial pressure; AIH, acute intermittent hypoxia; Control, time control without AIH; MSX-3, A2A receptor inhibitor; Ketanserin, 5-HT2A receptor inhibitor; Istradefylline, A2A receptor inhibitor; n’s, number of rats per group. All values are expressed at mean ± 1 SEM. Differences were considered significant if p < 0.050.

Table S2. Arterial PCO2, PO2, and MAP during hypoxic episodes

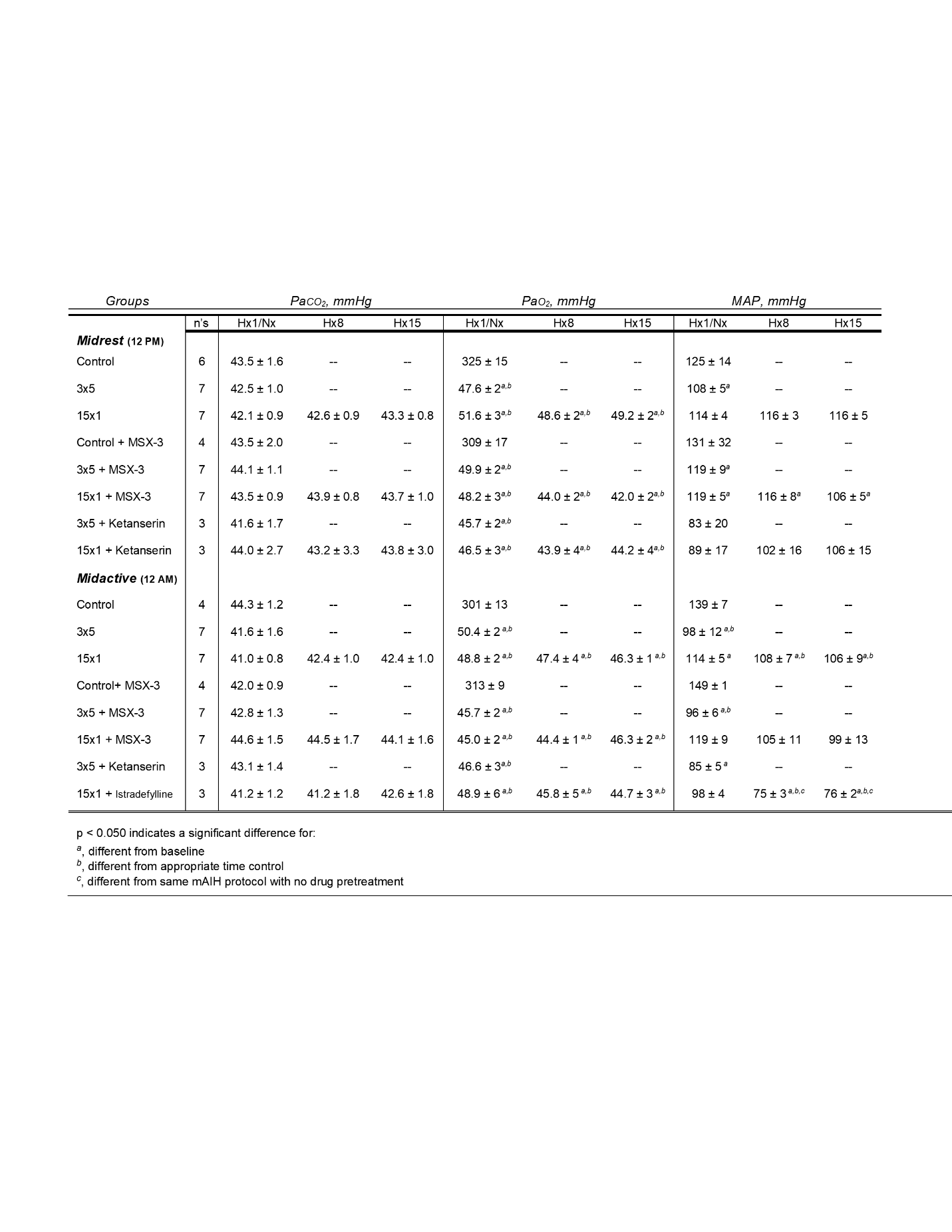

Hx1 designates hypoxic episode 1 for 3x5 and 15x1 mAIH protocols; Hx8 and Hx15 designate hypoxic episode 8 and 15 for the 15x1 mAIH protocol; Nx designates a blood gas sample taken in the same time period for time controls. A two-way mixed effects ANOVA followed by Tukey *post-hoc* test was utilized to determine effect of protocol with drug pretreatment on PaCO2, PaO2, and MAP within the midrest or midactive phase. MAP, mean arterial pressure; AIH, acute intermittent hypoxia; Control, time control without AIH; MSX-3, A2A receptor inhibitor; Ketanserin, 5-HT2A receptor inhibitor; Istradefylline, A2A receptor inhibitor; n’s, number for each group. All values are expressed as mean ± SEM. Differences were considered significant if p < 0.050.

Table S3. Phrenic nerve amplitude, in volts (V), at baseline and during maximum chemoreceptor stimulation

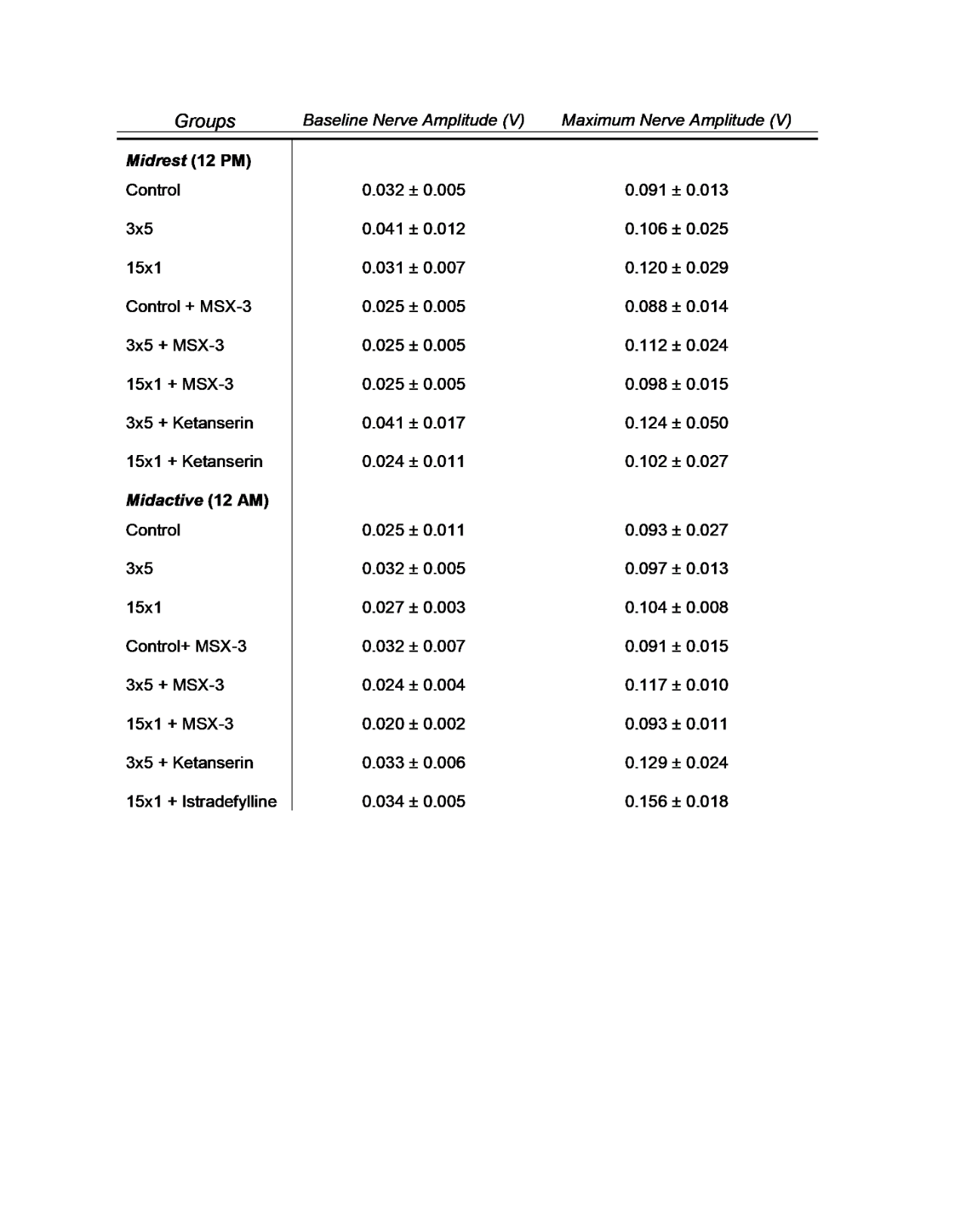

There was no effect of time of day or drug on baseline phrenic nerve amplitude, PNA (F_6,79_ = 1.075, p = 0.385; two-way ANOVA). There was no effect of time of day, drug pretreatment, or protocol on phrenic nerve amplitude during maximum chemoreceptor activation using 10% O2 and 7% CO2 with balanced N_2_ (F_15,70_ = 0.688, p = 0.788; three-way ANOVA). AIH, acute intermittent hypoxia; Control, time control without AIH; MSX-3, A2A receptor inhibitor; Ketanserin, 5-HT2A receptor inhibitor; Istradefylline, A2A receptor inhibitor. All values are expressed at mean ± SEM.

Table S4. Respiratory frequency (bpm) during baseline and 30, 60 and 90 min post-AIH

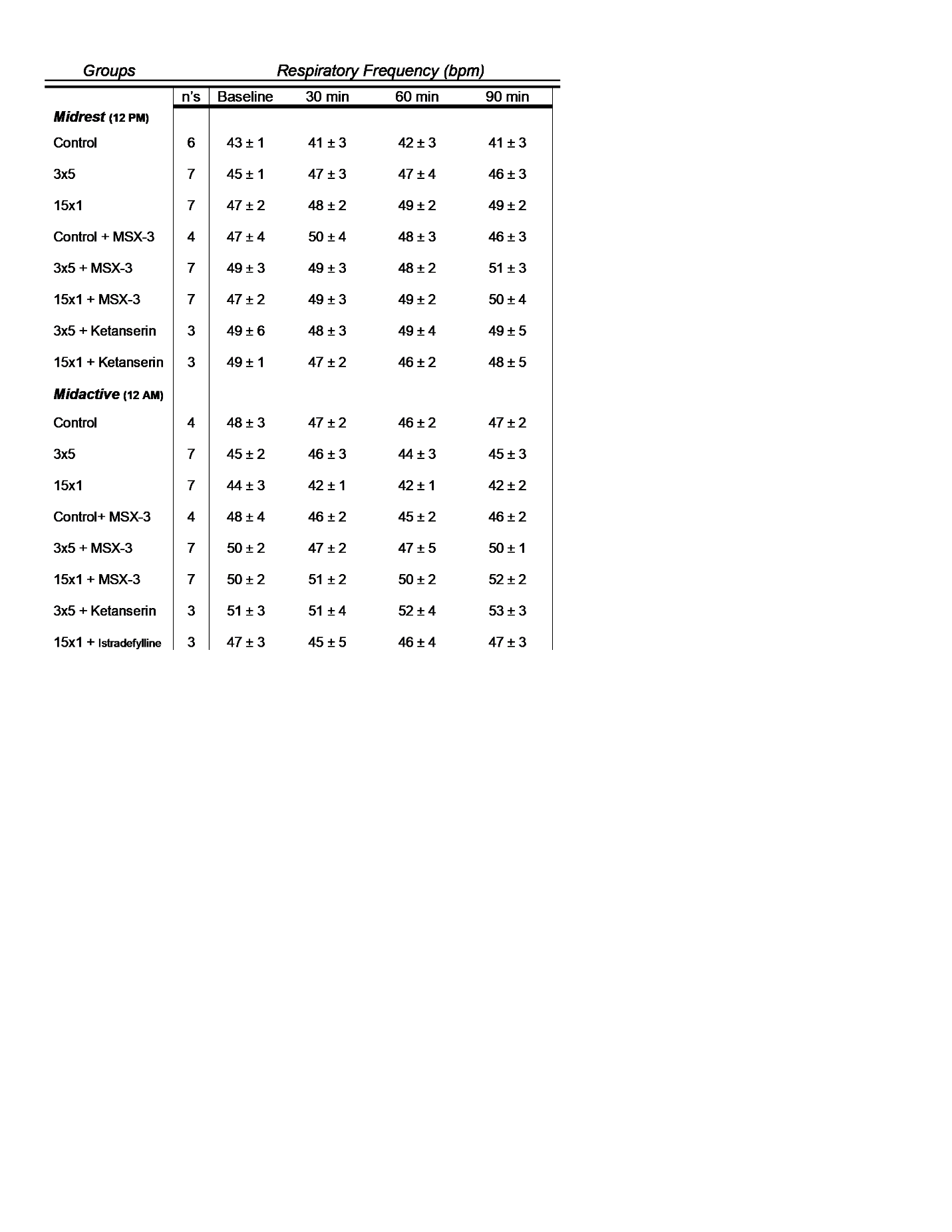

There was no effect of time of day or drug on baseline respiratory frequency (F_6,85_ = 1.154, p = 0.340; two-way ANOVA). There was no effect of protocol or drug pretreatment on respiratory frequency during midrest (F_21,108_ = 0.755, p = 0.767; three-way mixed effects ANOVA) or midactive (F_21,102_ = 0.457, p = 0.979; three-way mixed effects ANOVA) experiments. AIH, acute intermittent hypoxia; Control, time control without AIH; MSX-3, A2A receptor inhibitor; Ketanserin, 5-HT2A receptor inhibitor; Istradefylline, A2A receptor inhibitor; bpm, breaths per minute. All values are expressed at mean ± SEM.
